# Medulloblastoma Group 3 and 4 Tumors Comprise a Clinically and Biologically Significant Expression Continuum Reflecting Human Cerebellar Development

**DOI:** 10.1101/2022.06.09.495353

**Authors:** Daniel Williamson, Edward C Schwalbe, Debbie Hicks, Kimberly A Aldinger, Janet C Lindsey, Stephen Crosier, Stacey Richardson, Jack Goddard, Rebecca M Hill, Jemma Castle, Yura Grabovska, James Hacking, Barry Pizer, Stephen B Wharton, Thomas S Jacques, Abhijit Joshi, Simon Bailey, Steven C Clifford

## Abstract

Medulloblastoma is currently sub-classified into distinct DNA methylation subgroups/subtypes with particular clinico-molecular features. Using RNA-seq in large well annotated cohorts of medulloblastoma we show that transcriptionally Group3 and Group4 medulloblastomas exist not as discrete types but as intermediates on a bipolar continuum between archetypal Group3 and Group4 entities. Continuum position is prognostic, reflects propensity for specific DNA copy-number changes, key switches in isoform/enhancer usage and RNA-editing. Examining scRNA-seq profiles we show intra-tumoral transcriptional heterogeneity along the continuum is limited in a subtype-dependent manner. By integrating with a human scRNA-seq reference atlas we show this continuum is mirrored by an equivalent continuum of transcriptional cell types in early fetal cerebellar development. We identify unique developmental niches for all four major subgroups and link each to a common developmental antecedent. Our findings show a transcriptional continuum arising from oncogenic disruption of highly specific fetal cerebellar cell types, linked to almost every aspect of Group3/Group4 molecular biology and clinico-pathology.

## Introduction

The division of medulloblastoma into molecular subgroups has defined the past decade of medulloblastoma research making it all but impossible to interpret new findings except through the prism of these fundamental biological subdivisions. Medulloblastoma was first divided into subgroups on the basis of profiling by expression array (Cho et al., 2011; Fattet et al., 2009; Kool et al., 2008; Northcott et al., 2011; Thompson et al., 2006) and subsequently DNA methylation array (Hovestadt et al., 2014; Schwalbe et al., 2013). The current consensus is that there exist four major medulloblastoma subgroups (MB_SHH_, MB_WNT_, MB_Grp3_, MB_Grp4_) each with unique clinico-biological characteristics (Taylor et al., 2012); MB_WNT_ and MB_SHH_ are named after characteristic disruptions in the WNT (*CTNNB1* mutation (Ellison et al., 2005)) and SHH (*PTCH*, *SUFU*, *SMO* mutation or *GLI2* amplification (Kool et al., 2014)) pathways respectively. MB_WNT_ denotes an almost entirely curable disease (Ellison et al., 2005) and MB_SHH_ occur more frequently in infants (Kool et al., 2014). The remaining two subgroups Group3 (MB_Grp3_) and Group4 (MB_Grp4_) do not exhibit subgroup defining mutations (Northcott et al., 2017) but nonetheless possess distinct clinico-biological characteristics; MB_Grp3_ patients have a greater incidence of “high-risk” features such as LCA (large-cell/anaplastic) histology and *MYC* amplification (Kool et al., 2012; Northcott et al., 2012; Ryan et al., 2012; Taylor et al., 2012). MB_Grp4_ tumors more frequently demonstrate isochromosome 17q (i17q) (Sharma et al., 2019). The advent of routine medulloblastoma molecular subgrouping has enabled the current generation of molecularly driven trials (e.g. NCT02066220, NCT01878617, NCT02724579, NCT01125800) (Li et al., 2019; G. W. Robinson et al., 2015) which exploit MB_WNT_/MB_SHH_ biology to stratify treatments or direct biological therapeutics.

Further elaborations of the consensus subgroups were published, based primarily upon methylomic definitions (Cavalli et al., 2017; Northcott et al., 2017; Schwalbe et al., 2017). These were followed by a second consensus study which defined 8 subtypes within MB_Grp3_/MB_Grp4_ named I-VIII; a number of which comprised a mix of MB_Grp3_ and MB_Grp4_ tumors (Sharma et al., 2019). Furthermore, MB_SHH_ can be further divided into subtypes broadly associated with age at diagnosis^10,18,20^.

Based on murine modelling, expression and imaging studies (Gibson et al., 2010), MB_WNT_ and MB_SHH_ are believed to derive from two spatially distinct developmental origins in the early hindbrain; lower rhombic lip/dorsal brainstem and upper rhombic lip/early cerebellum, respectively. The developmental origins of MB_Grp3_ and MB_Grp4_ were investigated in a study mapping subgroup-specific super-enhancer elements, suggesting deep cerebellar nuclei residing in the nuclear transitory zone as the cell of origin for MB_Grp4_ (Lin et al., 2016). More recently, two studies which compared bulk and single-cell transcriptomic (scRNA-seq) MB profiles with developing murine cerebellar scRNA-seq reference datasets described MB_Grp3_ and MB_Grp4_ as most closely resembling Nestin positive stem cells (Vladoiu et al., 2019) and Unipolar Brush Cells (UBC) respectively, highlighting putative cells of origin (Hovestadt et al., 2019; Vladoiu et al., 2019). It is notable that the conclusions of each of these studies rely principally upon cross-species comparisons with murine as opposed to human developmental references. Human rhombic lip development is more complex and prolonged than that of mouse possessing unique features not shared with any other vertebrates (Haldipur et al., 2019).

Here, we characterize the transcriptomic landscape of 331 primary medulloblastoma, with clinico-pathological annotation, DNA methylation and copy number profiles, and we catalogue subgroup-specific isoforms and RNA-editing events. We show that, despite the discrete methylomic subdivisions of the MB_Grp3_/MB_Grp4_ methylation subtypes I-VIII, these tumors manifest transcriptionally on a bipolar continuum between MB_Grp3_ and MB_Grp4_ archetypes. Moreover, an individual tumor’s position on this continuum is predictive of methylation subtype, prognosis, specific copy number and mutational alterations, and activation of key molecular pathways and regulatory events. By using human scRNA-seq fetal cerebellar reference data, we show that this continuum mirrors and recapitulates the major developmental trajectories within early human cerebellar development allowing us to map the interplay between key oncogenic events and putative cells of origin for each medulloblastoma subtype.

## Results

### Medulloblastoma shows a continuum of expression between MB_Grp3_ and MB_Grp4_

RNA-seq (∼90M paired-end reads) was performed on 331 snap-frozen primary samples from patients with a diagnosis of medulloblastoma (Supplementary Table 1). Transformed gene-level read counts were subject to consensus NMF clustering with resampling to determine the most stable number of clusters and metagenes i.e. major biological effects described by multiple genes and summarized as a single score. As expected, a 4-metagene/4-cluster solution was optimally stable, reflecting the four major consensus subgroups as currently understood (Figure 1A). ∼3% (10/331) of samples were defined as non-classifiable i.e. low probability of classification. ∼4% (13/331) samples could only be classified as indeterminate MB_Grp3_/MB_Grp4,_ i.e. confidently classifiable as either MB_Grp3_ or MB_Grp4_ but not specific as to which. The distribution of clinico-biological features was consistent with previously described features of the consensus MB subgroups (Figure 1A & 1SA); for instance, Chromosome 6 loss in 83% (24/29) of MB_WNT_.

**Figure 1.**
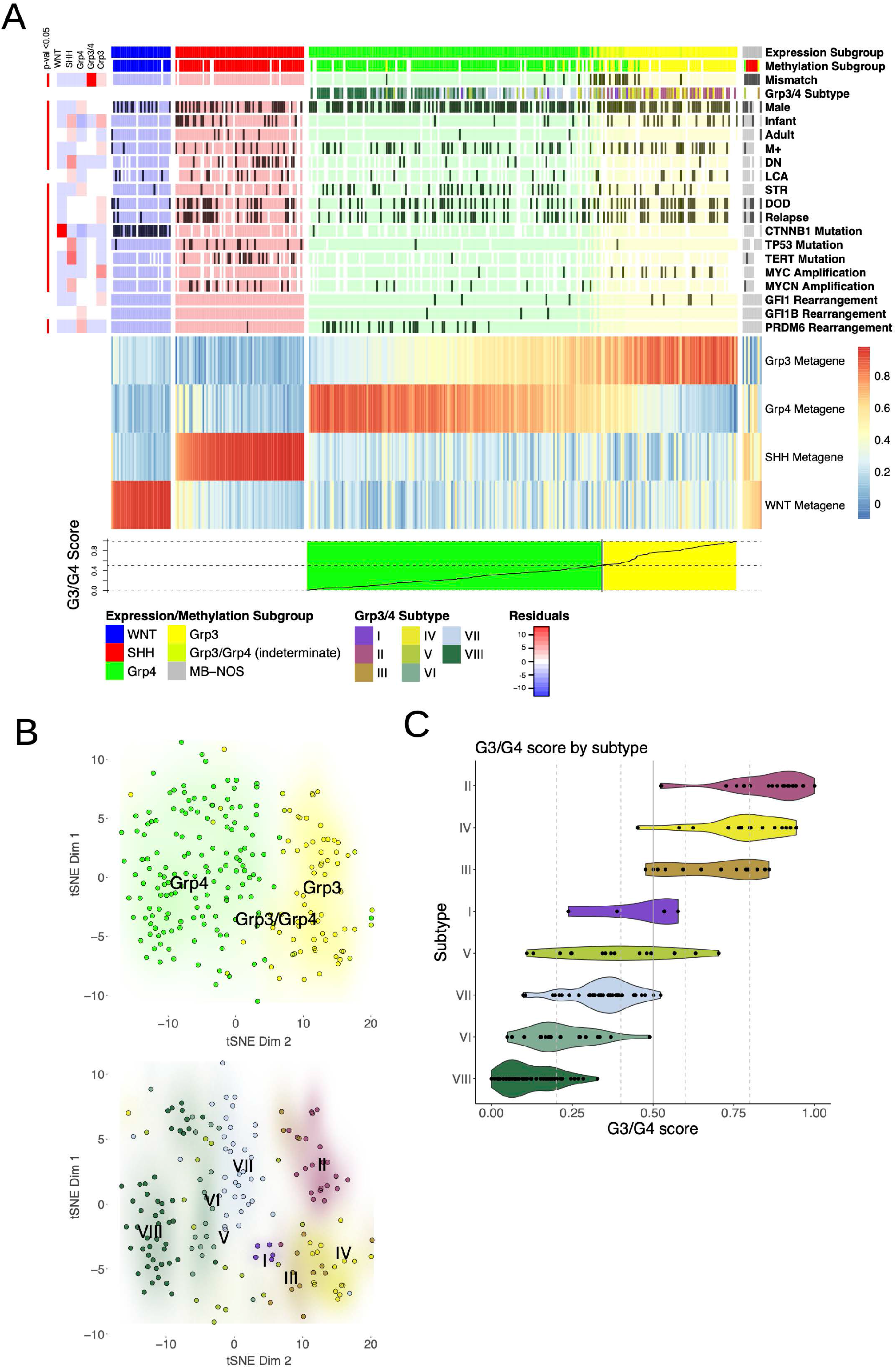
**A:** Heatmap showing 4 consensus NMF metagenes calculated for n=331 MB and grouped by subgroup. MB_Grp3_/MB_Grp4_ individuals are ordered by G3/G4 score. Annotation shows subgroup as determined by RNA-seq (Expression Subgroup), subgroup as determined by methylation (Methylation Subgroup), methylation MB_Grp3_/MB_Grp4_ subtype (I-VIII) as per Sharma *et al* 2019(Sharma et al., 2019) defined using MNPv2 classifier(Capper et al., 2018) (Grp3/4 Subtype). All other characteristics are indicated to be present or not by dark grey shading according to the following scheme: Infant = age at diagnosis < 3 years, Adult = age at diagnosis > 16 years, DN = Desmoplastic/Nodular, LCA = Large-cell/anaplastic, STR = Sub-Total Resection, DOD=Dead of Disease. Side annotation (top left) shows a heatmap of chi-square residuals indicating subgroup enrichment and significance where relevant. The line plot (bottom) shows G3/G4 score. **B:** t-SNE plot showing MB_Grp3_/MB_Grp4_ samples shaded by subgroup (top) and methylation MB_Grp3_/MB_Grp4_ subtype (I-VIII) (bottom). **C:** Violin plot showing G3/G4 score by MB_Grp3_/MB_Grp4_ subtype (I-VIII).

The two metagenes which described MB_Grp3_ and MB_Grp4_ samples were notably gradated and overlapping in an anti-correlative manner (Figure 1A) implying that, contrary to some previous descriptions using expression microarrays (Cavalli et al., 2017), MB_Grp3_ and MB_Grp4_ are not distinct transcriptional entities but rather exist as a continuum between two transcriptional polarities we refer to here as G3 and G4. To describe this continuum, we created a continuous score (G3/G4-score) scaled between 0 and 1 to reflect the proportionate amount of G3/G4 metagene expression in each MB_Grp3_/MB_Grp4_ (i.e. all non-WNT/non-SHH tumors) whereby a score of ‘0’ indicates a 100% G4 tumor and ‘1’ 100% G3 (Figure 1A). This was applied to the 223 samples classified as MB_Grp3_, MB_Grp4_ or intermediate MB_Grp3_/MB_Grp4_.

For convenient comparison, we sub-divided the expression continuum (G3/G4-score) into five notional groups: HighG4 (0-0.2, n=69/223 (31%)), LowG4 (0.2-0.4, n=60/223 (27%), G3.5 (0.4-0.6, n=39/223 (17%)), LowG3 (0.6-0.8, n=22/223 (10%)) & HighG3 (0.8-1 G3/G4-score, n=33/223 (15%)). All samples with >0.5 G3/G4 score were classified as MB_Grp3._ Notably, 15/20 (75%) MB_Grp3_/MB_Grp4_ samples which showed disagreement in classification between RNA-seq and DNA methylation array were classified as indeterminate MB_Grp3_/MB_Grp4_ by RNA-seq (Figure 1A). Examining the MB_Grp3_/MB_Grp4_ subtype (I-VIII) calls by t-SNE (Figure 1B) shows clustering by subtype, suggesting that each methylation subtype imparts distinct secondary expression characteristics beyond the primary G3/G4 continuum metagene. Regardless, the MB_Grp3_/MB_Grp4_ subtypes may be broadly ordered upon the G3/G4 continuum in partially overlapping domains from most Group4-like to most Group3-like (VIII, VI, VII, V, I, III, IV, II respectively) (Figure 1C).

Specific clinico-biological features were significantly non-randomly distributed across the G3/G4 continuum (Figure 2A). For instance, LCA pathology is significantly enriched at the G3 end of the continuum (D=0.339, p=0.046, n=158); MB_Grp3_ patients display LCA (Large Cell Anaplasia) pathology ∼3 times more frequently if HighG3 as opposed to LowG3. Large (arm level/chromosomal) copy number alterations are likewise non-randomly distributed with respect to the G3/G4 continuum. Most notably, i17q is proportionately greater in HighG4 individuals (75% compared to 36% in Low G4, D=0.402, p<0.001, n = 201) and chromosome 8 gain more frequent in HighG3 (44% in High G3, 5% in Low G3, D=0.69, p<0.001, n=201) (Figure 1S). Mutations are not frequent in MB_Grp3_/MB_Grp4_ (Northcott et al., 2017), however non-synonymous mutations of *ZMYM3*, *KDM6A* are significantly non-randomly distributed with respect to the continuum (each p<0.01)(Figure 2S).

**Figure 2.**
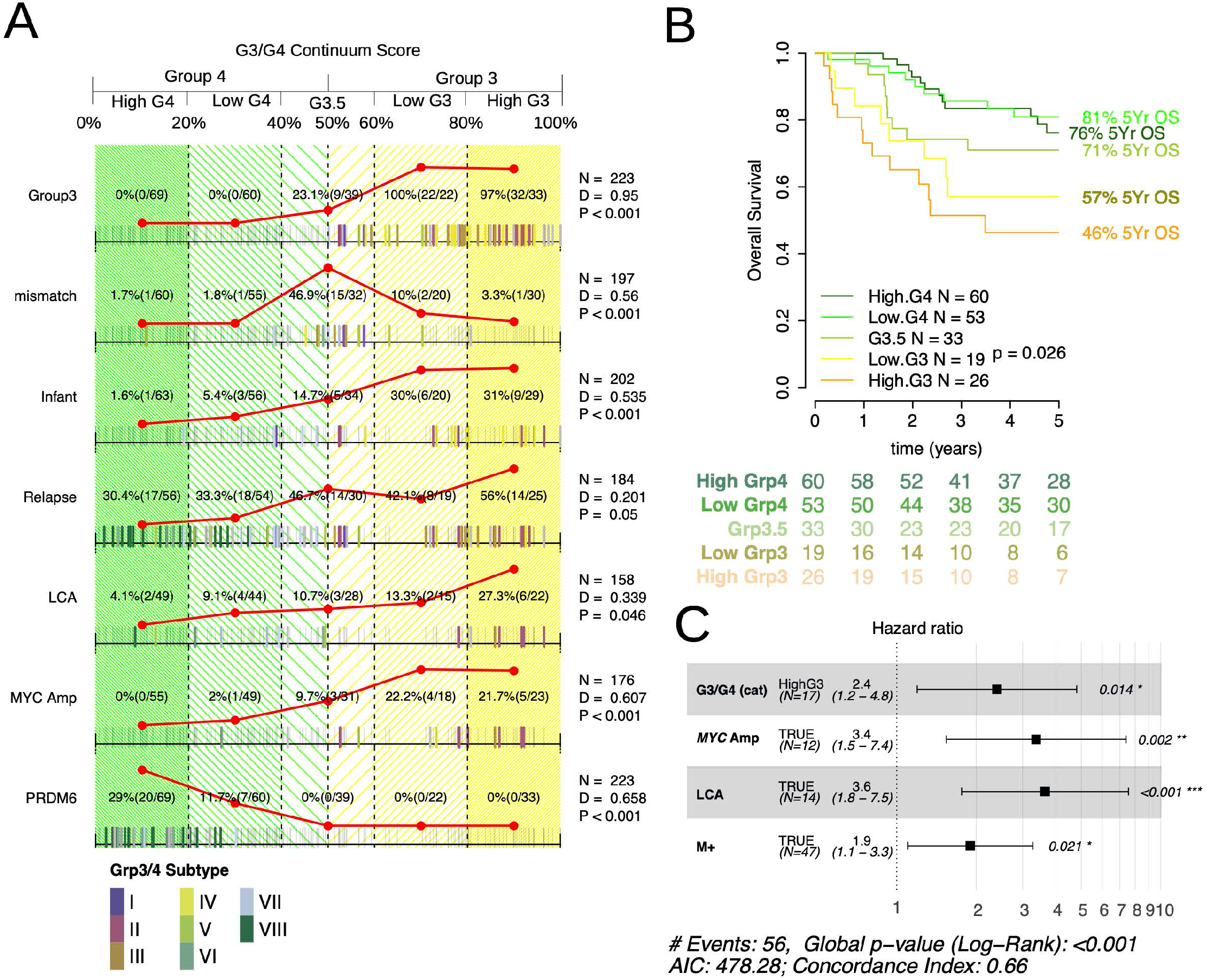
**A:** Rug plot showing distribution of clinico-path features with respect to G3/G4 score. Summary counts are given according to the divisions of HighG4, LowG4, G3.5, LowG3, HighG3 and reflected by the red line plots. Presence of a feature is indicated by a bold tick mark the color of which indicates methylation MB_Grp3_/MB_Grp4_ subtype (I-VIII). Adjusted p-values for a Kolmogorov-Smirnoff statistic (D) are shown to denote non-random distribution of features with respect to G3/G4 score. Mismatch=mismatch between methylation and expression call, Infant=age at diagnosis < 3 years, M+=Metastatic, DOD=Dead of Disease, LCA = Large-Cell/anaplastic, PRDM6 = *PRDM6* rearrangement. **B:** Kaplan-Meier plot showing significant differences in MB_Grp3_/MB_Grp4_ overall survival by G3/G4 continuum position. **C:** Forest plot showing a multivariate Cox model fitted to progression free survival and containing the independently significant variables High G3, *MYC* amplification, LCA and M+.

We examined the relationship between the G3/G4 score and prognosis. Treating the G3/G4 score as categorical showed a progressively poorer 5-year overall survival (OS) across the continuum; Log-rank p=0.026, n=191, HighG3=46%, LowG3=57%, G3.5=71%, LowG4=81%, HighG4=76% (Figure 2B). Cox-regression indicates the continuous G3/G4 score is highly significant (p=0.003, n=191) showing an increase in relative risk of death (RR) of 4.7 times greater for a patient with a G3/G4 score 0 compared to a score of 1. Using the G3/G4 score as a categorical variable, only HighG3 patients have significantly poorer prognosis (RR=2.8, p=0.001, n=191), Multivariable Cox-regression analysis of progression free survival including other risk factors *MYC* amplification, LCA histology and metastatic disease showed that the HighG3 category retains significance (RR=2.4, p=0.014, n=135) indicating the G3/G4 score possesses significant independent prognostic power distinct from its association with other “high-risk” disease features (Figure 2C).

### A G3/G4 continuum score can be reverse-engineered from DNA methylation profiles to validate clinico-pathological associations

A series of sample cohorts of MB_Grp3_/MB_Grp4_ with DNA methylation profiles have previously been published by ourselves and others (Cavalli et al., 2017; Northcott et al., 2017; Schwalbe et al., 2017; Sharma et al., 2019). To these we added 166 profiles to produce a large cohort (n=1670) better powered to validate and further expand our findings made using transcriptomic datasets. We therefore explored the possibility of reverse-engineering a G3/G4 score from DNA methylation data. Using the same method as used for expression was impossible, given that the constrained range (i.e. 0 (fully unmethylated) to 1 (fully methylated)) and bimodal distribution of CpG methylation does not lend itself straightforwardly to a continuous score (Figure 3SA). Unlike expression, which tends to follow a log-linear association with G3/G4 score, methylation follows a sigmoidal distribution from hypo- to hyper-methylation or *vice versa*. The inflection point along the G3/G4 continuum at which these CpGs “switch” from one state to the other varies by CpG (Figure 3A, 3SB,C). We trained a classifier using a training cohort of MB_Grp3_/MB_Grp4_ samples for which we possessed both RNA-seq and DNA methylation profiles (n=192). Pre-selecting 400 cross-validated CpG features which distinguish between each of the G3/G4 categorical states we used these to train a random forest classifier to accurately predict (RMSE = 0.036) a G3/G4 score from DNA methylation data alone (Figure 3B).

**Figure 3.**
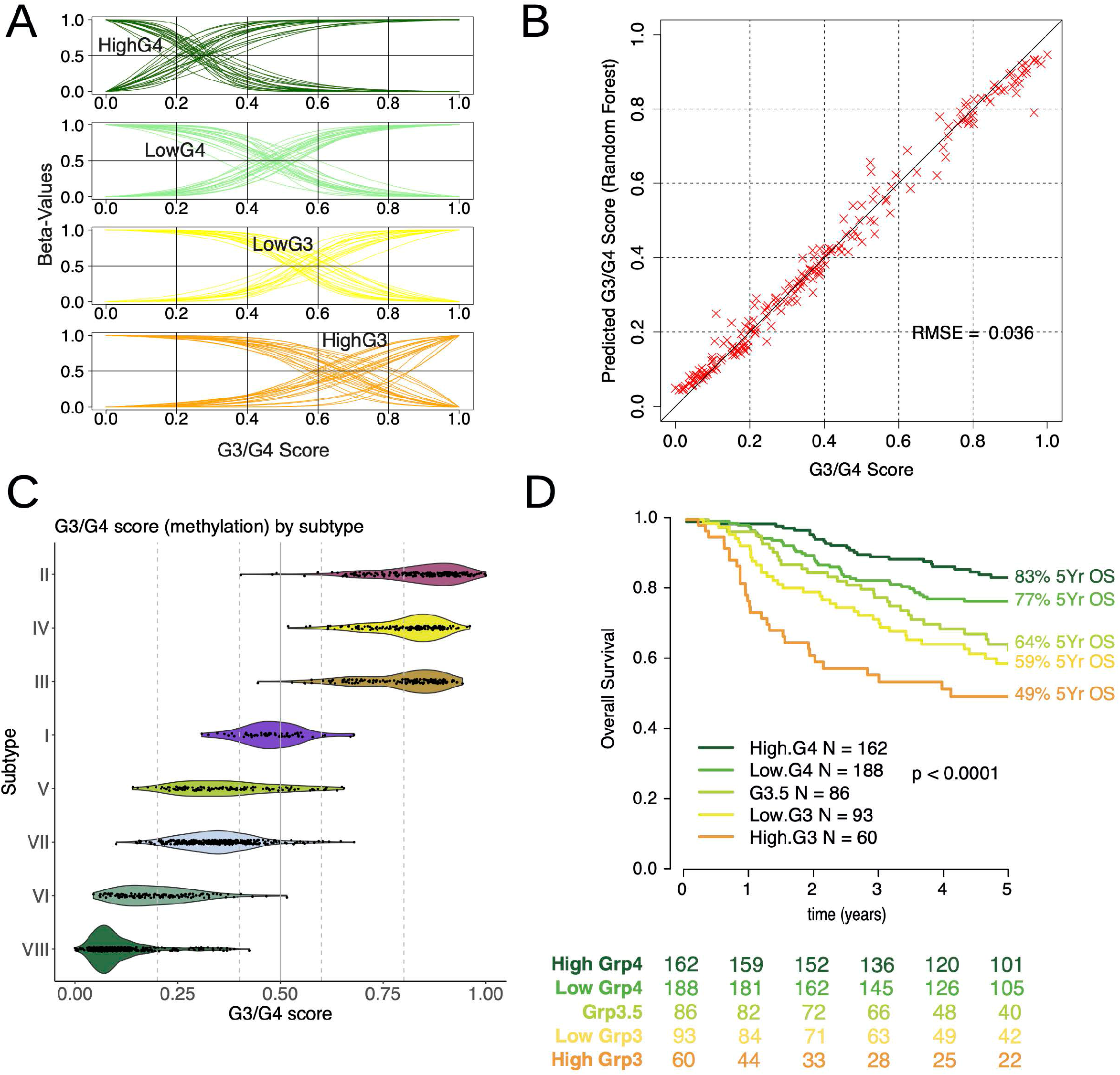
**A:** Fitted sigmoid curve representing the relationship between CpG beta-value and G3/G4 Score. Top 40 most discriminatory CpGs distinguishing HighG4 (dark green), LowG4 (light green), Low G3 (yellow) and High G3 (orange) are shown. **B:** The performance of the cross-validated random forest classifier showing predicted G3/G4 score (derived from DNA methylation values) against actual G3/G4 score (derived from RNA-seq) n = 192. **C:** Violin plot showing G3/G4 score (derived from methylation) by MB_Grp3_/MB_Grp4_ (I-VIII) subtype. **D:** Kaplan-Meier plot showing significant differences in MB_Grp3_/MB_Grp4_ overall survival in patients aged > 3 years by G3/G4 score (as derived from methylation values) n = 589.

Using this larger MB_Grp3_/MB_Grp4_ methylation cohort we were able to demonstrate significant non-random distribution with respect to the continuum of infant status (<3 years), metastases, LCA, and *MYC* amplification (each more frequent in HighG3 patients) and mutations of *PRDM6*, *KDM6A*, *KMT2C* and *ZMYM3* (more frequent in HighG4 patients) (each p<0.001, Figure 4SA). Likewise, chromosomal gains of 1q, 5, 6, 8 and 16q (each p<0.001 and most frequent in HighG3) and i17q (p<0.001 most frequent in HighG4) (Figure 4SA). These findings thus validated our findings from the initial RNA-seq cohort.

The larger cohort size allowed us to also explore the relationship between the G3/G4 continuum and the MB_Grp3_/MB_Grp4_ subtypes (I-VIII) as well as their previously reported clinico-pathological/mutational characteristics (Sharma et al., 2019). The MB_Grp3_/MB_Grp4_ subtypes as predicted from DNA methylation data once again occupy discrete but partly overlapping domains within the G3/G4 continuum; broadly ordered - as per the RNA-seq only cohort - from most archetypally MB_Grp4_ to MB_Grp3_ VIII, VI, VII, V, I, III, IV, II (Figure 3C).

We next asked whether the variation in distribution of clinicopathological features and mutation previously described as being characteristic of MB_Grp3_/MB_Grp4_ subtypes (I-VIII) (Sharma et al., 2019) were attributable to position on the G3/G4 continuum, MB_Grp3_/MB_Grp4_ subtype (I-VIII) or indeed both. Certain frequent clinicopathological features and copy number changes (e.g. metastatic disease, *MYC* amplification, LCA histology, i17q, loss of chromosome 8, gain of chromosome 5) are significantly non-randomly distributed with respect to G3/G4 continuum, even within individual subtypes (Figure 4SB,C). For example, 100% (11/11) of subtype III with *MYC* amplifications are HighG3 compared to 59% (69/117) without *MYC* amplification. The presence of i17q as the only major chromosomal alteration is a highly characteristic change in subtype VIII but when considering only medulloblastoma subtype VIII is still significantly enriched at the High G4 end of the continuum (D=0.162, p=0.014). The relative contribution of MB_Grp3_/MB_Grp4_ subtype and G3/G4 continuum to predicting presence/absence of a clinicopathological or mutational change was additionally demonstrated by logistic regression (Figure 4SD). This showed that, in several instances (e.g. LCA, *MYC* amplification, i17q as only chromosomal aberration) the response variable was better described using the G3/G4 continuum rather than the medulloblastoma subtype (I-VIII) as an explanatory variable.

The relationship between G3/G4 score and risk of death is significant and striking, allowing us to validate the findings of our RNA-seq cohort with greater confidence; patients age >3 years Log-rank p<0.0001 n=589, HighG3=49%, LowG3=59%, G3.5=64%, LowG4=77%, HighG4=83% (Figure 3D, 5SA). A similar result is found in patients of all ages; Log-rank p<0.0001 n=654 (Figure 5SA). Modelling G3/G4 score as a continuous variable using a Cox proportional hazards again in patients age > 3years shows a 3x increased risk of death from one end of the continuum to the other (RR=3, n=589, p<0.001) (Figure 5SB). MB_Grp3_/MB_Grp4_ subtypes (I-VIII) are also significantly associated with overall survival (n=524, p<0.001) (Figure 5SC).

### The G3/G4 continuum is associated with differential regulation of oncogenic/developmental pathways

The expression of 590 genes are significantly correlated with the G3/G4 score in our RNA-seq cohort (p<0.01, log_2_ fold change>10, n=223), increasing/decreasing log-linearly across the continuum. Most notably, *MYC* expression correlates significantly with G3/G4 score (Rho=0.73, p<0.001, n = 223). HighG3 samples had a mean expression of *MYC* 46 times greater than HighG4 and 2 times greater than LowG3 (Figure 4A). Performing Gene Set Enrichment Analysis (GSEA) we observed that transcriptional targets of MYC were also significantly upregulated (NES = 3.37, p=0.007) (Figure 4B). ssGSEA analysis (Hänzelmann et al., 2013) was used to represent activation/repression of pathway/signatures for each individual and found several oncogenic pathways which were progressively activated or repressed in a manner significantly correlated (each p<0.001) with the G3/G4 continuum including MYC, Cell Cycle, MTOR, TGF-Beta (activated at the G3 pole) and NOTCH (activated at the G4 pole) (Figure 4C). In addition, a broad pattern of progressive neuronal differentiation at the G4 pole and photoreceptor (CRX/NRL) characteristics at the G3 pole of the G3/G4 continuum were observed.

**Figure 4.**
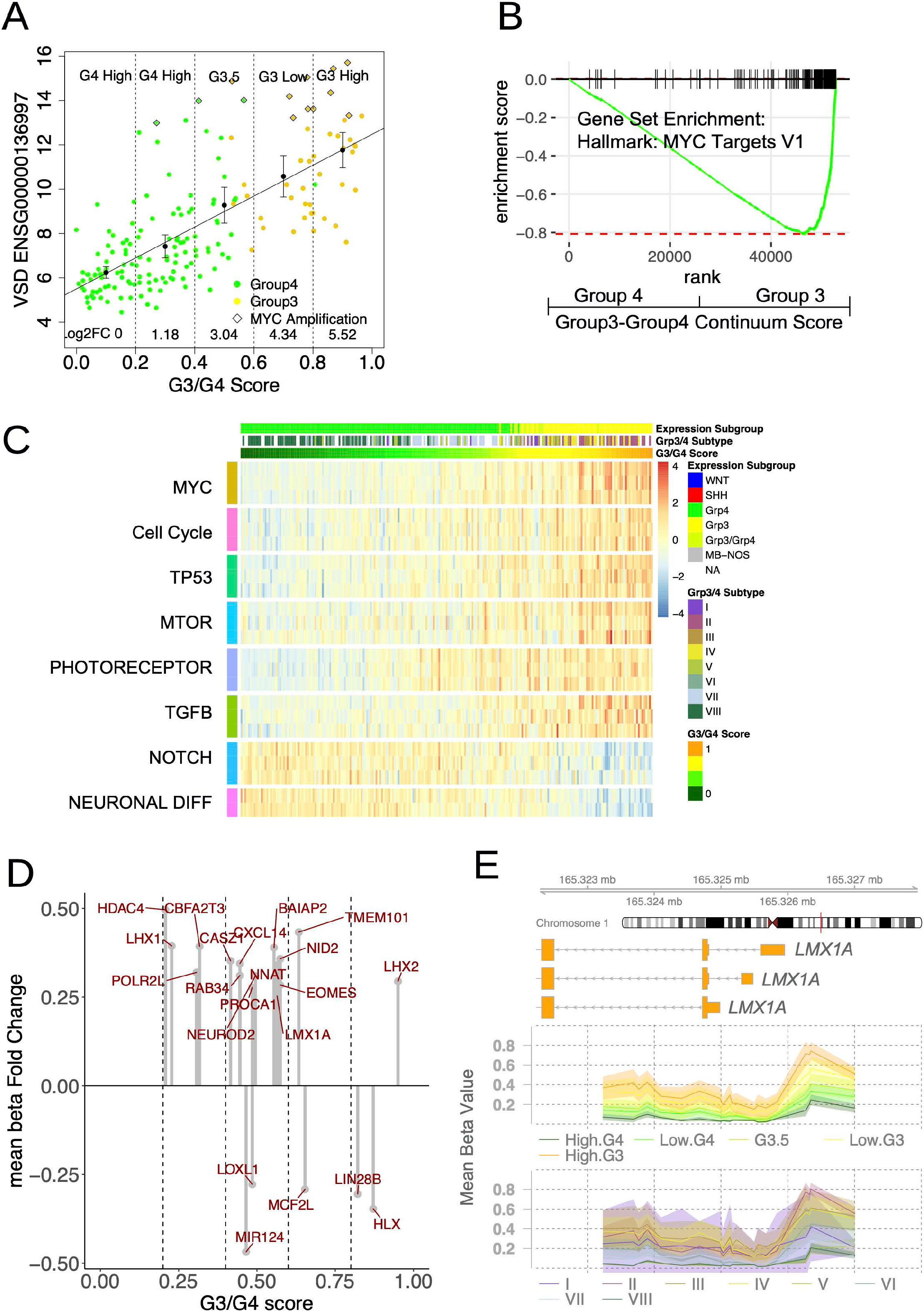
**A:** Scatterplot showing significant correlation (p<0.001) between *MYC* expression and G3/G4 score. Log-linear line of best fit is shown. Dotted lines divide into High G4, Low G4, G3.5, Low G3, High G3 and Log2 fold change for each category relative to HighG4 are shown. **B:** GSEA enrichment plot showing significant enrichment of MYC target genes. Genes were ranked by correlation with G3/G4 score. **C:** Heatmap of ssGSEA results showing level of pathway enrichment for 223 MB_Grp3_/MB_Grp4_ individuals ordered by G3/G4 score. MsigDB pathways are curated into pathways (see methods). **D:** Lollipop plot showing mean beta fold change for DMRs within MB_Grp3_/MB_Grp4_ specific enhancers/superenhancers. The position on the x axis reflects the average point on the continuum at which the methylation level switches from hypo- to hypermethylation. **E:** Plot showing an MB_Grp3_/MB_Grp4_ specific enhancer within the MB_Grp3_ specific gene *LMX1A* which overlaps with a differentially methylated region significantly associated with the G3/G4 continuum. The mean beta value per G3/G4 category (High G4, Low G4, G3.5, Low G3, High G3) and MB_Grp3_/MB_Grp4_ subtype (I-VIII) is shown by line and the 95%CI by shaded area.

We examined differentially methylated regions (DMRs) within previously identified MB_Grp3_/MB_Grp4_ specific enhancer loci(Lin et al., 2016), identifying 45 which also overlapped with gene promoters; each “switched” from hypomethylated to hypermethylated or *vice versa* at specific points along the G3/G4 continuum. The expression of 33/45 of these genes are significantly correlated with the G3/G4 continuum (p<0.01). This switching appears progressive, with certain MB_Grp3_/MB_Grp4_ enhancer loci “switching” earlier and others later. For instance, the enhancer/DMR loci overlapping with the promoters of medulloblastoma lineage development/differentiation genes *LHX1*, *NEUROD2*, *LMX1A*, and *HLX* on average “switch” at points 0.23, 0.49, 0.56, 0.87 on the G3/G4 continuum (Figure 4D,E). We note also that expression of each of these genes is significantly correlated with the G3/G4 continuum and DMR methylation (p<0.01). This is consistent with a developmental identity controlled by cumulative changes in underlying epigenetic architecture driving progressive transition from an MB_Grp3_ to a MB_Grp4_ cell state.

### The G3/G4 continuum is associated with post-transcriptional regulation of isoform expression and RNA-editing

To explore the clinico-biological significance of differentially expressed transcriptional isoforms across subgroups, Kallisto (Bray et al., 2016) was used to estimate abundance. Taking TPM (Transcripts Per Million) >10 as indicative of a moderate-highly expressed isoform it is notable that the diversity of isoforms being expressed across subgroups was significantly greater in MB_Grp4_ than MB_Grp3_ (p<0.001, F=9.877) (Figure 6SA). 153 genes were identified whose expression overall is invariant but for which expression of specific isoforms correlates significantly with G3/G4 score (Figure 5A). For instance, overall expression of *GTF2I (General Transcription Factor IIi)* is ubiquitous but a progressive isoform switch corresponding to the balance between β/δ (*GTF2I-215*/*GTF2I-218*) and α/γ (*GTF2I-221*/*GTF2I-212*) isoforms correlates significantly to G3/G4 score (Figure 5B). These isoforms switches are known to alter protein stability (Shirai et al., 2015) and subcellular localization (Shirai et al., 2017).

**Figure 5.**
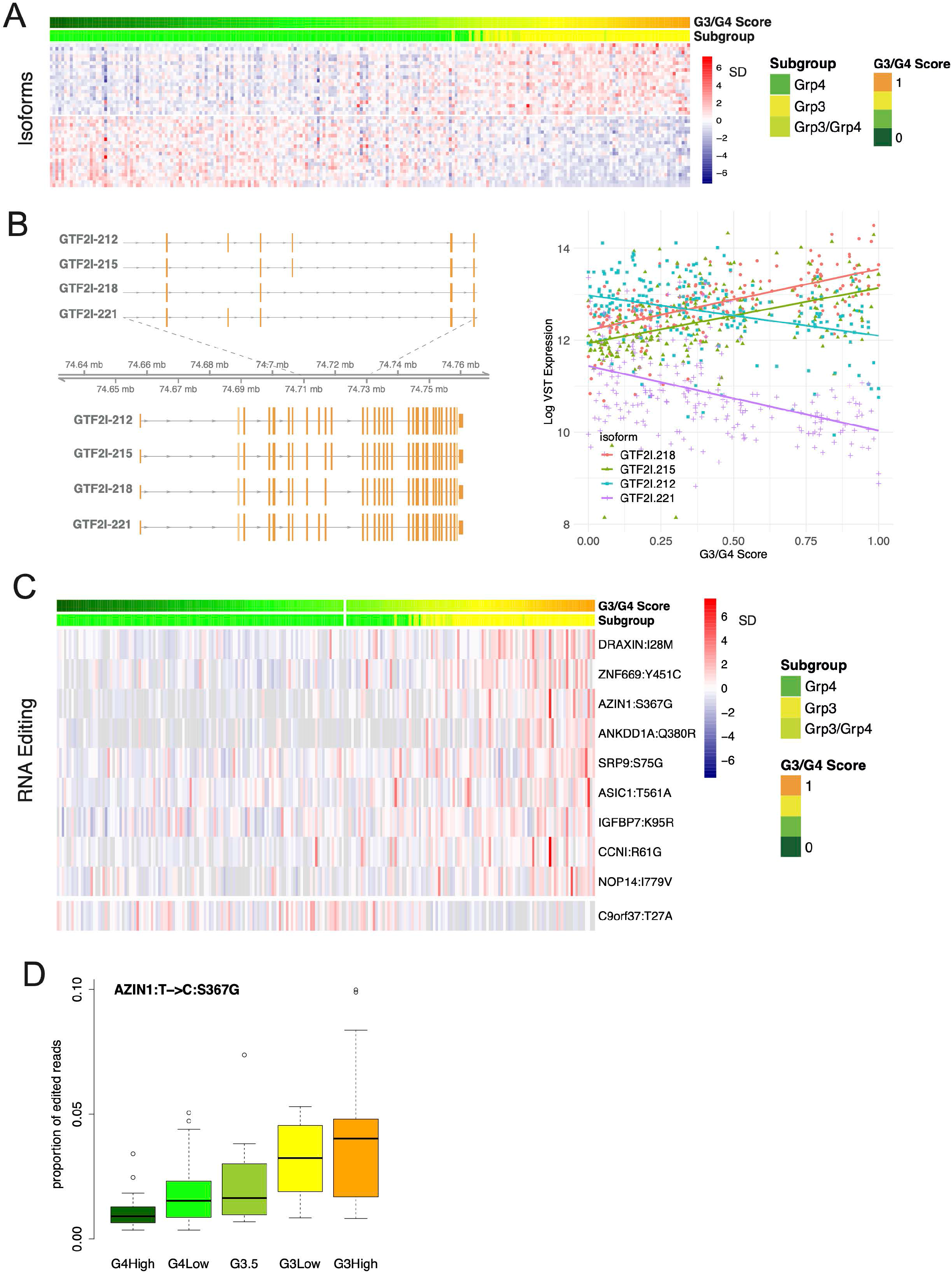
**A:** Heatmap showing expression of top significantly differentially expressed isoforms of genes whose overall expression is otherwise not significantly differentially expressed with respect to G3/G4 score. **B:** Schematic showing exon structure of 4 *GTF2I* isoforms significantly differentially expressed with respect to G3/G4 score (left) and scatterplot showing expression of these *GTF2I* isoforms vs G3/G4 score; line represents fitted log-linear model NB: *GTF2I* is not significantly differentially expressed at the gene level. **C:** Top 10 significantly differentially edited non-synonymous RNA editing positions with respect to G3/G4 score. **D:** Boxplot showing level of T>C RNA editing at a non-synonymous position S367G within *AZIN1;* level of editing is significantly associated with G3/G4 score.

4,668,508 established RNA editing sites were profiled using the QEdit/Reditools pipeline (Giudice et al., 2020); we observed significant differences in overall A-I editing level. The OEI (Overall Editing Index - i.e. total number of reads with G at all known editing positions over the number of all reads covering the positions) differs significantly with respect to subgroup (F=9.761, n=223, p<0.001); post-hoc testing showed RNA editing events in MB_Grp4_ to be significantly more numerous than MB_Grp3_ and MB_SHH_ (each p<0.01) (Figure 6SB). Analysis of 5174 non-synonymous RNA editing sites showed 32 significantly differentially edited with respect to the G3/G4 continuum (p<0.05, Figure 5C). One such RNA editing site is *AZIN1* chr8:103841636T>C, known to result in a S367G substitution which causes conformational changes, cytoplasmic-to-nuclear translocation and gain of function increasing tumor potential in hepatocellular carcinoma (Chen et al., 2013), non-small-cell lung cancer (Hu et al., 2017), colorectal cancer (Shigeyasu et al., 2018) and gastric cancer (Okugawa et al., 2018) (Figure 5D). It is also notable that *ADAR1* and *ADAR2* expression are both correlated with G3/G4 score (rho=0.54, p<0.001 and rho=0.33, p<0.001, n=223, respectively).

### Intra-tumoral cellular heterogeneity with respect to the G3/G4 continuum is apparent but constrained by subtype

We projected our MB_Grp3_/MB_Grp4_ metagenes onto a MB_Grp3_/MB_Grp4_ scRNA-seq dataset comprising 4,256 cells from 15 individuals (5xSubtype-II, 2xSubtype-III, 1xSubtype-I, 2xSubtype-V, 4xSubtype-VIII) previously published by Hovestadt *et al* (Hovestadt et al., 2019). This allowed us to estimate a G3/G4 score for each cell within a given sample. MB_Grp3_ individuals were described by Hovestadt *et al* as being dominated by cells with an undifferentiated progenitor-like expression program and MB_Grp4_ dominated by a differentiated neuronal-like program; these appear to broadly equate with our MB_Grp3_ and MB_Grp4_ metagenes. MB_Grp3_/MB_Grp4_ intermediate tumors were described by Hovestadt *et al* as a partial admixture of cell types. The distribution of G3/G4 scores at the single cell level shows that, whilst there is a certain amount of intra-tumoral cellular variation (Figure 6A), the majority of cells fall within the same G3/G4 range observed in the equivalent subtype bulk RNA-seq profiles (Figure 6B). For example, amongst medulloblastoma subtype VIII individuals 78% (667/853) of cells fall within the HighG4 range, 22% (184/853) within LowG4 and 0.2% (2/853) within G3.5. This is equivalent to the bulk profiles for which 83% (40/48) of individuals fall within the HighG4 range and 17% (8/48) fall within the LowG4 range (Figure 6B). In short, the phenomenon of a G3/G4 continuum observed in bulk RNA-seq analysis seems not to be produced by differently proportioned admixtures of otherwise wholly Group3 and Group4 cells but rather medulloblastoma are composed of populations of individual cells, which themselves display continuous G3/G4 expression characteristics; these being constrained to occupy a discrete part of the G3/G4 continuum as dictated by their MB_Grp3_/MB_Grp4_ (I-VIII) subtype.

**Figure 6.**
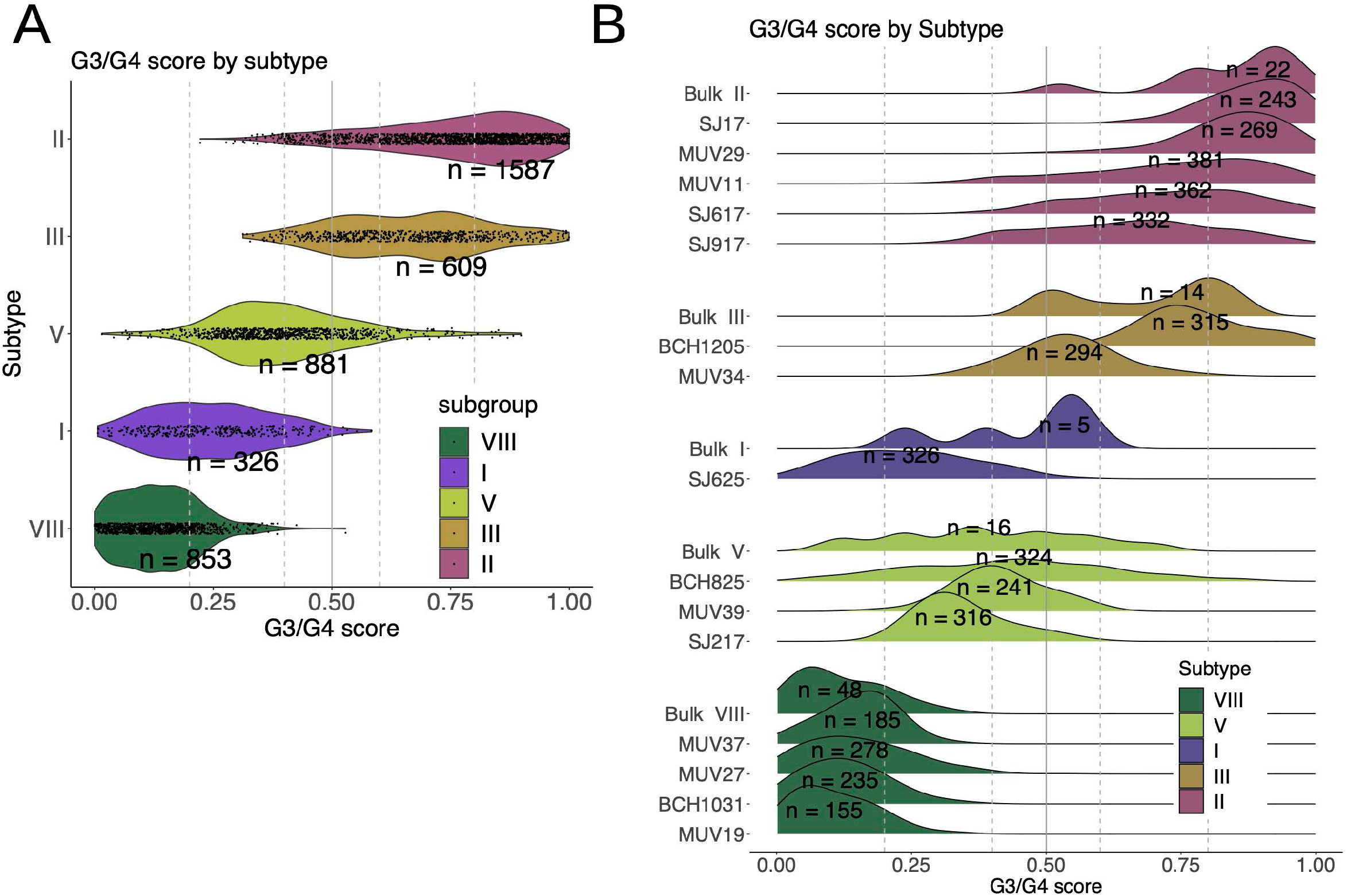
**A:** Violin plot showing per-cell G3/G4 score (derived from projection onto scRNA-seq data) for 15 MB_Grp3_/MB_Grp4_ patients aggregated by subtype. **B:** Ridgeplot showing distribution of per-cell G3/G4 score (derived from projection onto scRNA-seq data) for each of 15 MB_Grp3_/MB_Grp4_ patients shown alongside the G3/G4 score distribution of equivalent subtype bulk tumors. n=x refers to number of individuals for bulk tumors and number of cells for the scRNA-seq data.

### Medulloblastoma subtypes and the G3/G4 continuum are mirrored in early human cerebellar development

The origins of medulloblastoma within spatially and temporally distinct regions of the fetal cerebellum (Upper Rhombic Lip/Granule Cell lineage for MB_SHH_ and Lower Rhombic Lip for MB_WNT_) have been primarily established by mouse modelling (Gibson et al., 2010; Lin et al., 2016) and, more recently, by comparison with reference to mouse fetal cerebellum scRNA-seq datasets which suggest a unipolar brush cell (UBC) origin for MB_Grp4_ (Hovestadt et al., 2019; Vladoiu et al., 2019). Such comparisons in embryonal tumors are predicated on the idea that partial transformation in an early prenatal cell interrupts development/differentiation resulting in a proportion of the expression characteristics of the tumor initiating cell being retained.

Here, we avoid any cross-species comparisons by using instead a human fetal cerebellum scRNA-seq reference set (69,174 cerebellar cells 9-21 post conception weeks (PCW)). We reconstructed a pseudotemporal cellular trajectory within a broadly defined rhombic lip lineage (12,243 cells, comprising rhombic lip precursors (RL), excitatory cerebellar nuclei/unipolar brush cells (eCN/UBC), granule cell precursors (GCP), and granule cell neurons sub-divided into four clusters (GN)) (Figure 7A). We projected our 4 subgroup metagenes onto these cerebellar cells, identifying those cells which showed highest expression of each metagene. These cells occupy distinct branches of our lineage. High MB_WNT_ metagene expressing cells, as expected, occupy a discrete subset of the RL precursors (Figure 7B). High MB_Grp3_/MB_Grp4_ metagene expressing cells occupy a distinct eCN/UBC branch beginning with RL precursors (highly expressing MB_Grp3_ metagenes) and transitioning midway to eCN/UBC cells highly expressing the MB_Grp4_ metagene (Figure 7B); this cell trajectory in effect mirrors the G3/G4 continuum. This can be demonstrated formally by calculating a projected per-cell G3/G4 score, revealing a smooth transition from a MB_Grp3_-like to a MB_Grp4_-like expression state (Figure 7C). More straightforwardly, this is demonstrated by observing the significant change in expression with respect to pseudotime of those G3/G4 continuum-associated genes whose expression is sufficiently high to be consistently detectable within the relatively low depth scRNA-seq data (each p<0.01, Figure 7SA).

**Figure 7.**
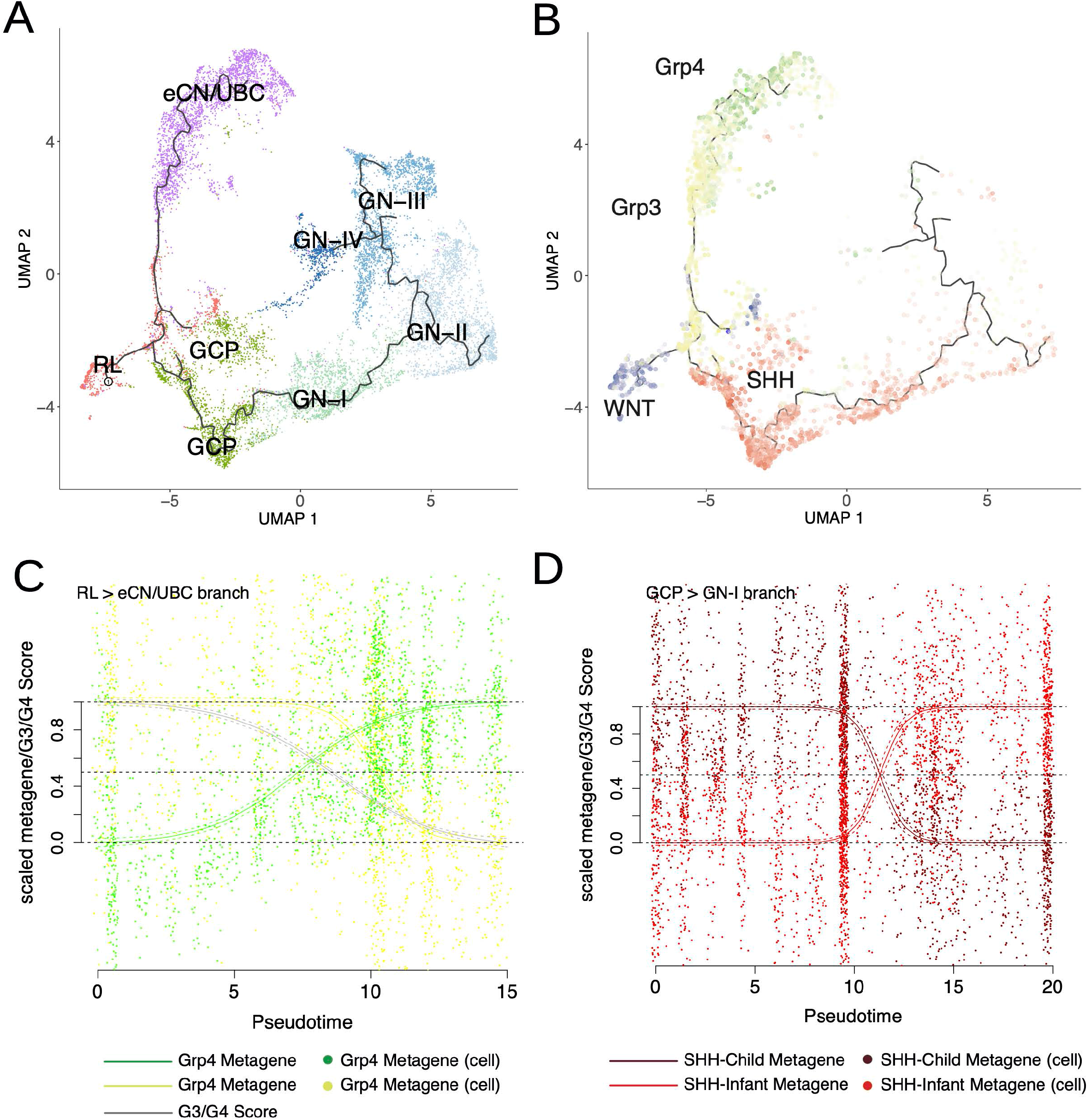
**A:** UMAP plot of scRNA-seq profiles showing 12,243 cells of the rhombic lip lineage arranged according to developmental trajectory which is indicated by black line. Color denotes cell type as determined by graph based clustering; RL=Rhombic Lip precursors, GCP=Granule Cell Precursors, GN-I, GN-II, GN-III, GN-IV = four Granule Neuron cell types, eCN/UBC=excitory Cerebellar Neurons/Unipolar Brush Cells. **B:** UMAP plot of the rhombic lip lineage with those cells within the top decile of metagene expression are marked with the following colors: MB_Grp4_=green, MB_Grp3_=yellow, MB_SHH_=red, MB_WNT_=blue. **C:** Scatterplot showing per-cell scaled metagene expression along the RL to eCN/UBC branch. Fitted sigmoid curves are shown with SD indicated as dashed lines. The grey line represents a sigmoid curve fitted to per-cell G3/G4 score as a function of pseudotime. **D:** Scatterplot showing per-cell scaled metagene expression along the GCP to GN branch. Fitted curves are shown with SD shown as dashed lines. Curves are scaled to be constrained to a range of 0 and 1, in order to be coherent with bulk analysis. For this reason, by definition, some individual cells lie outside the 0 and 1 range.

Cells which express the MB_SHH_ metagene most highly, as expected, occupy a granule cell developmental branch beginning with GCPs and extending partly into the earliest GN cell types (Figure 7B). Two metagenes representing MB_SHH-Infant_ (primarily patients <4 years) and MB_SHH-Child_ (primarily patients > 4 years) - as described in previous studies (Kool et al., 2014; Schwalbe et al., 2017) - were also projected onto the cells in this branch. This indicated a switch midway through the granule cell pseudotemporal lineage from a predominantly MB_SHH-Infant_ metagene to a predominantly MB_SHH-Child_ metagene expression; this coincided approximately with the first transition from GCPs to GNs (Figure 7D). Again, where expression of individual genes which distinguish infant MB_SHH_ from childhood MB_SHH_ were sufficiently detectable within the scRNA-seq profiles they were significantly associated with pseudotime (each p<0.01, Figure 7SB).

Thus, by aligning the oncogenic G3/G4 scale with the pseudotemporal scale we were able to order and assign tumorigenic events to specific points within fetal cerebellar developmental lineages (Figure 8). *MYC* amplification, for instance, tends to coincide with the earlier RL pseudotemporal space as opposed to *KDM6A* mutation which occupies the later more differentiated eCN/UBC space. Likewise for aneuploidies, gain of chromosome 8 coincides with the earlier RL developmental space and i17q (as the sole copy number alteration) with the later eCN/UBC cell types.

**Figure 8.**
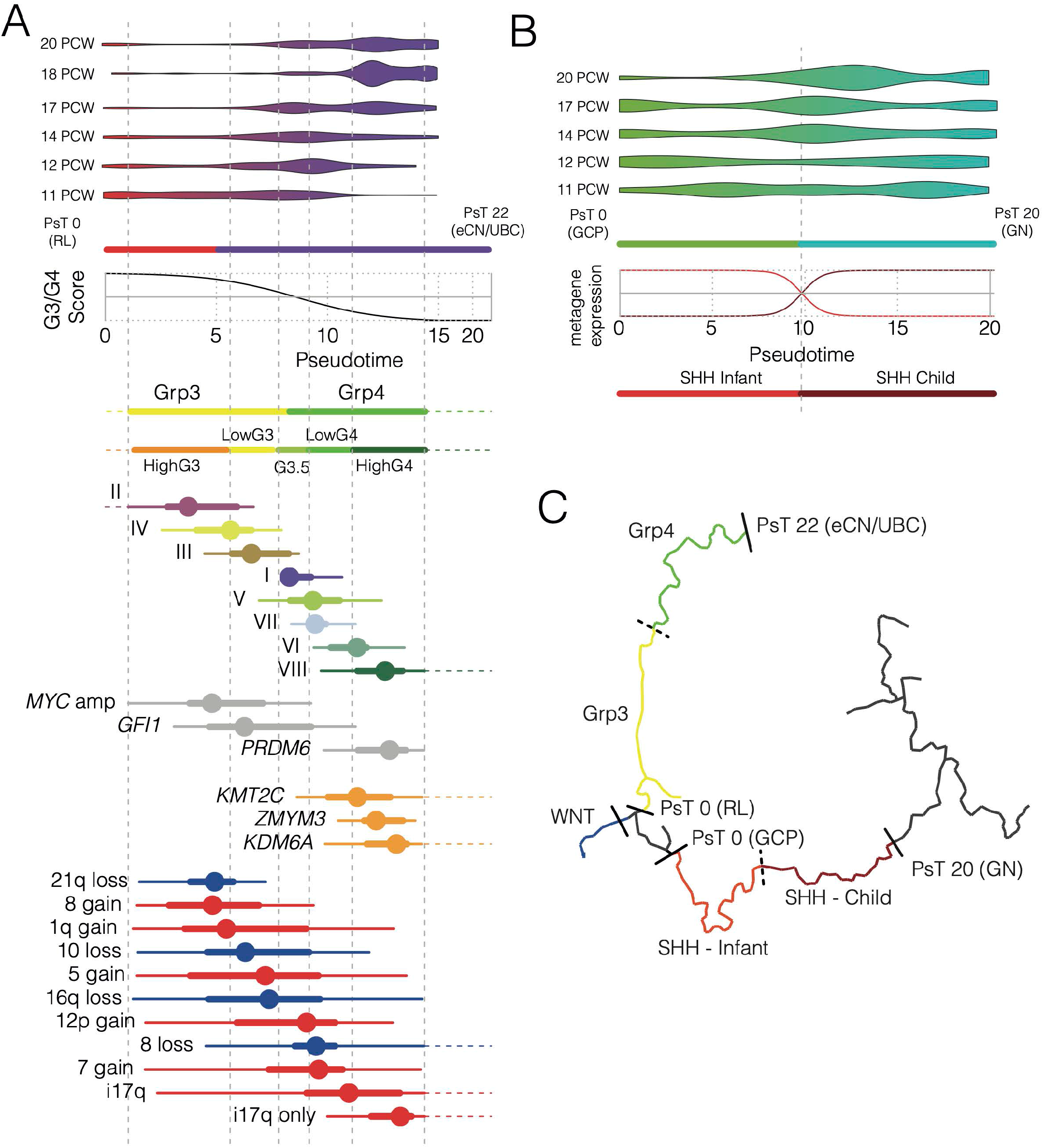
**A:** Schema showing the RL to eCN/UBC developmental branch, the relationship between pseudotime and G3/G4 score and the staging of key tumor characteristics. From top to bottom: a violin plot showing pseudotime distribution of cells by time of sampling, color transition red to purple marks the point along the developmental trajectory where cells are defined as eCN/UBC. A fitted sigmoid curve showing the relationship between pseudotime and G3/G4 score. Tumor characteristics are transformed from the G3/G4 scale to the pseudotime scale and marked at the appropriate points. Color bars represent subgroups. Methylation subtypes (I-VIII), mutations and copy number changes are marked by box and whisker. Dot represents median distribution thick line represent the interquartile range and the thinner lines corresponds to range. Dotted lines denote where the range extends up to a G3/G4 score of 0 and 1, i.e. matching the *ne plus ultra* pseudotime after which G3/G4 score is unchanged and exact relationship must be extrapolated. **B:** Schematic showing the GCP to GN developmental branch, the relationship between pseudotime and MB_SHH-Infant_ or MB_SHH-Child_ metagene. From top to bottom: a violin plot showing pseudotime distribution of cells by time of sample, color transition green to blue marks the point along the developmental trajectory where cells become defined as GN. A loess curve showing the relationship between pseudotime and MB_SHH-Infant_ (red) or MB_SHH-Child_ metagene (dark red). Color bars show parts of trajectory paralleled by MB_SHH-Infant_ or MB_SHH-Child_ tumors. **C:** UMAP of developmental trajectory marked with colors to denote parts most associated with each MB subgroup and the relevant pseudotime (PsT) scale.

We note that as well as the pseudotemporal transition from MB_Grp3_ to MB_Grp4_ or MB_SHH-Infant_ to MB_SHH-Child_ there is also a literal temporal transition as well. The cerebellar cells most closely associated with the archetypal MB_Grp3_ are predominant at 11PCW (and possibly before). By 18PCW, those most closely associated with the archetypal MB_Grp4_ predominate; this persists until at least 20PCW and between 12PCW and 14PCW, cells associated with G3.5 and LowG4 predominate. This temporal staging from early to late forms of MB_Grp3_/MB_Grp4_ is also mirrored in the average age of onset of disease. The distribution of age at diagnosis of each MB_Grp3_/MB_Grp4_ (I-VIII) subtype closely parallels the distribution across the G3/G4 continuum (Figure 8SA) and there is a significant correlation between G3/G4 score and age at diagnosis (Figure 8SB). On the RL to GN branch, cells most closely associated with MB_SHH-Infant_ are predominant at PCW11 and reduced by PCW20 at which point MB_SHH-Child_ associated cells predominate (Figure 8B).

We also note that the relative frequencies of MB_Grp3,_ MB_Grp4_ and the distribution of the patient population across the G3/G4 continuum is mirrored in the size of the relevant pseudotemporal space; by space we mean the estimated proportion of putative cells of origin for each subtype (adjusted by number of cells sampled) as a function of the length of time they persist throughout development (9-21 PCW) (Figure 8SC,D). For instance, in our methylation cohort the incidence of MB_Grp3_ relative to MB_Grp4_ is approximately 1:2 (35% 585/1670 MB_Grp3_ vs 65% 1085/1670). The overlap with the relevant proportion of matched pseudotemporal space indicates the same 1:2 ratio (38% MB_Grp3_ matched space vs 62% MB_Grp4_ matched space). The bimodal empirical distribution of the G3/G4 continuum is also paralleled by the pseudotemporal space occupied (Figure 8SD). We regard the above observations as likely explanatory of the relative differences in age of onset and frequency of the different MB_Grp3_/MB_Grp4_ subtypes and reflective of the relative window of opportunity in terms of time and developmental space for transformation to occur.

## Discussion

Here we show that, with regards to their transcriptomes, the primary inter-tumoral variation in MB_Grp3_/MB_Grp4_ patients is continuous, in contrast to the discrete nature of the methylation MB_Grp3_/MB_Grp4_ subtypes (I-VIII) (Cavalli et al., 2017; Northcott et al., 2017; Schwalbe et al., 2013; Sharma et al., 2019). This is not in itself contradictory, as we show that the MB_Grp3_/MB_Grp4_ methylation subtypes are ordered along the G3/G4 continuum in discrete but partially overlapping domains (Figure 1D). Furthermore, as has been demonstrated previously (Cavalli et al., 2017; Sharma et al., 2019), the methylation subtypes are reflected to some extent in their expression profiles (Figure 1C). Nonetheless, these are shown here to be secondary expression characteristics subordinate to the overarching primary expression characteristic which is the G3/G4 continuum.

The position of an individual MB_Grp3_/MB_Grp4_ tumor upon the continuum is significantly associated with certain mutations, copy number aberrations, clinico- and histopathology. This is to be expected, as many of these have been shown to be non-randomly associated with MB_Grp3_/MB_Grp4_ subtypes (Sharma et al., 2019). That both methylation subtype and the expression continuum are related to key tumor characteristics and indeed to one another is clear. The question remains to what extent the inter-tumoral variation in such characteristics may be better explained by position upon the continuum than by methylation subtype. For at least some of these characteristics - those which are frequent and not specific to single subtypes (e.g. *MYC* amplification, LCA, i17q, gain of chromosome 5 and loss of chromosome 8) - the answer appears to be that they are better explained by the continuum (Figure 4SB, C, D).

The most striking association is between the G3/G4 continuum and risk of death; at least during the first 5 years post-diagnosis. Risk increases continuously with the G3/G4 continuum (Figure 3D), the documented phenomenon (Sharma et al., 2019) of late (>5-year post diagnosis) relapse in subtype VIII notwithstanding. We regard this study as a description of an extremely close and therefore important relationship between biology and clinical course rather than an advocation for its use as a clinical biomarker. Those judgements should be made using prospective clinical trials and the cohort used here, whilst sizeable and carefully reviewed, is a retrospective cohort with all the limitations and caveats that implies. Nevertheless, we note that when it comes to incorporating molecular data into risk stratification schemes the use of a single G3/G4 risk score for all MB_Grp3_/MB_Grp4_ patients has a certain pragmatic logic over atomizing a rare cancer into 8 separate subtypes.

Pathway analysis of the G3/G4 continuum shows a concomitant activation of oncogenic processes (e.g. MYC, MTOR, TP53) as tumors become more MB_Grp3_-like, which itself suggests a more aggressive phenotype. The influence of the G3/G4 continuum also extends to post-transcriptional regulation i.e. isoform usage and RNA-editing. A close relationship with cell differentiation (e.g. CRX/NRL, neuronal differentiation) is also evident and consistent with previous descriptions of MB_Grp3_/MB_Grp4_ biology cell identity and differentiation (Bandopadhayay et al., 2019; Garancher et al., 2018). This is further reflected in the progressive switches in methylation status we observe within MB_Grp3_/MB_Grp4_ specific enhancers (Lin et al., 2016).

Importantly, we show here that the MB_Grp3_/MB_Grp4_ continuum is not produced as the result of an admixture of discrete cell types in different proportions but by individual cells which themselves exist upon the same expression continuum as the bulk tumors. In part, this was observed by Hovestadt *et al* (Hovestadt et al., 2019) in their original analysis of their pooled MB_Grp3_/MB_Grp4_ scRNA-seq data. They described two metagenes diverging according to *MYC* expression and describe bulk tumors as composed of cells of either a predominately MB_Grp3_, MB_Grp4_ or intermediate type which themselves represent a continuum of neuronal differentiation (Hovestadt et al., 2019). We have expanded this by fitting individual cells onto the same metagene scale used to define the bulk tumor transcriptome, thereby defining more precisely the range of transcriptional intra-tumoral heterogeneity within MB_Grp3_/MB_Grp4_ tumors and showing that it appears to be confined to certain limits prescribed by MB_Grp3_/MB_Grp4_ subtype. This is in turn consistent with the finding that medulloblastoma sampled from different areas of the tumor or at diagnosis and relapse rarely alter subgroup (Kumar et al., 2021; Morrissy et al., 2016; Ramaswamy et al., 2013).

Unlike previous studies which attempted to define cells of origin we used a human rather than a mouse scRNA-seq reference set for comparison. The use of a human atlas is significant because human RL persists longer through cerebellar development than the mouse and has unique cytoarchitectural features not shared with any other vertebrates (Haldipur et al., 2019). Mouse RL is a transient, proliferative stem cell zone present between E12.5 and E17.5 whereas human RL begins as a progenitor niche and is later compartmentalized into ventricular and subventricular zones, forming a human-specific progenitor pool within the posterior lobule, which persists until birth (Haldipur et al., 2019). We show that the MB_Grp3_/MB_Grp4_ continuum is paralleled by a fetal cerebellar lineage that begins with an RL progenitor and ends with eCN/UBC. This points to highly specific windows of opportunity - which we define in part here - within developmental pseudotemporal space for oncogenic events to successfully provoke medulloblastoma of a given subtype. These windows of opportunity are highly coherent with both the age of onset of the disease and the relative frequency of the individual subtypes. Importantly, we were able to identify a developmental niche for each of the 4 main medulloblastoma subgroups including a separate space for MB_SHH-Child_ and MB_SHH-Infant_. Each of these is contained within a branch of the same early cerebellar lineage explicitly unifying each of the 4 subgroups to a common developmental antecedent; something not reported in previous studies. For instance, Hovestadt *et al* (Hovestadt et al., 2019) were unable to identify a significant matching reference cell type for MB_Grp3_ and MB_WNT_ whereas Vladoiu *et al* (Vladoiu et al., 2019) did not analyze MB_WNT_ and note a prosaic resemblance of MB_Grp3_ to Nestin^+^ early neural stem-like cells. We should note that we were selective, albeit based on prior knowledge, in the subset of cell types we considered as potential candidate cells of origin: figuratively by assigning them to what we broadly described as the Rhombic lip lineage and literally by the physical process of cell extraction and the points in early human development for which sampling was possible (PCW 9-21). MB_WNT_ in particular is thought to originate in the dorsal brainstem, and it may be that certain alternative cells of origin were excluded or curtailed on that basis. Nevertheless, previous studies follow a similar logic to our own and the coherent picture of the relationships between the subgroups would seem to bear out our choices.

In conclusion, our findings point to the following important insights. First, that Group3/Group4 MB and their methylation subtypes exist transcriptionally upon a continuum and that this is mirrored entirely by an equivalent continuum of transcriptional cell types in early human fetal cerebellar development. Second, that by using a human scRNA-seq reference all four MB subtypes can be linked to a common developmental antecedent within the RL lineage. Third, that transcriptional intra-tumoral heterogeneity is limited to certain domains within the continuum as dictated by subtype and finally, that the continuum is linked with almost every aspect of Group3/Group4 molecular biology and clinico-pathology. We anticipate this to have implications for the future treatment and modelling of the disease, most pressingly a need to match cell type with specific timing of mutations to develop faithful models.

## Acknowledgements

Supporting Grants: This work was supported by Cancer Research UK (C8464/A13457 & C8464/A2339), Tom Graham Trust/CCLG and by the INSTINCT network, co-funded by The Brain Tumour Charity, Great Ormond Street Children’s Charity, and Children with Cancer UK (grant 16/193). TSJ is grateful for additional funding from the Olivia Hodson Cancer Fund, Cancer Research UK and the National Institute of Health Research. All research at Great Ormond Street Hospital NHS Foundation Trust and UCL Great Ormond Street Institute of Child Health is made possible by the NIHR Great Ormond Street Hospital Biomedical Research Centre. The views expressed are those of the author(s) and not necessarily those of the NHS, the NIHR or the Department of Health.

## Author Contributions

Conceptualization D.W., E.C.S., S.B. and S.C.C; Methodology D.W., E.C.S., S.B. and S.C.C; Formal Analysis D.W., E.C.S, K.A.A. and Y.G.; Investigation D.W., E.C.S, D.H, K.A.A, J.C.L, S.C., S.R., J.G., R.M.H., J.C., S.B.W., T.S.J., A.J., S.B. and S.C.C.; Resources K.A.A., B.P., S.B.W., T.S.J. and A.J.; Data Curation D.W., E.C.S., D.H., J.C.L., S.R., J.G., R.M.H., J.C., Y.G., and J.H.; Writing – Original Draft D.W., E.C.S and S.C.C; Writing – Review & Editing D.H., K.A.A., S.R., J.G., R.M.H, B.P., and S.B.; Visualization D.W. and J.H.; Supervision D.W., S.B. and S.C.C.; Funding Acquisition D.W., S.B. and S.C.C. All authors approved final version of the manuscript.

## STAR Methods

### Resource Availability

#### Lead contact

Further information and request for resources and reagents should be directed to and will be fulfilled by the lead contact, Daniel Williamson.

#### Materials availability

This study did not generate new unique reagents.

#### Data and code availability

Data arising from this study has been deposited in E-MTAB-10754 and E-MTAB-10767 (ArrayExpress) and are publicly available as of the date of publication. Additionally, this study makes use of previously deposited data sets GSE130051, GSE93646, GSE119926 (GEO/NCBI). For scRNA-seq fetal cerebellar data, processed data are available through the Human Cell Atlas (https://www.covid19cellatlas.org/aldinger20) and the UCSC Cell Browser (https://cbl-dev.cells.ucsc.edu). Sequence data is available in the Database of Genotypes and Phenotypes, under accession number phs001908.v2.p1 (dbGAP/NCBI). Details are listed in the key resources table and method details section. No custom code was used in this study. Open-source algorithms were used as detailed in the method details section. Details on how these algorithms were used are available from the corresponding authors upon request.

### Experimental Models

#### Human tissue samples

Snap frozen tumor samples from individuals with confirmed medulloblastoma diagnosis were used for RNA-seq analysis. These were provided as part of UK CCLG-approved biological study BS-2007–04 and/or with approval from Newcastle North Tyneside Research Ethics Committee (study reference 07/Q0905/71); informed, written consent was obtained from parents of all patients younger than 16 years.

### Method Details

#### Patient samples and study cohort

331 tumor samples from individuals with confirmed medulloblastoma diagnosis were used for the RNA-seq analysis. Histopathological variants were defined according to the WHO 2016 guidelines (Louis et al., 2016). Metastatic status (M+) was defined as M>1 as per Chang’s criteria (Chang et al., 1969). *MYC* and *MYCN* amplification status was assessed by fluorescence in situ hybridization and/or copy-number estimates from methylation array and *TP53*, *CTNNB1*, and *TERT* mutation status by Sanger sequencing. DNA was extracted using Qiagen DNeasy blood and tissue kit. Other mutations were assessed using next-generation sequencing. Whole-exome and targeted gene panel sequencing was performed using the Agilent SureSelect target enrichment platform and Illumina paired-end sequencing according to manufacturer’s instructions. NGS datasets were analyzed for coding/exonic region variants using Genome Analysis Toolkit (GATK) version 3.7, according to Broad Institute’s best practices (Burrows wheeler alignment, Haplotype Caller, Variant Quality Score Recalibration for exomes and Hard-filtering for panel) (Van der Auwera et al., 2013) and annotated using Ensembl Variant Effect Predictor (McLaren et al., 2016). Variants were predicted pathogenic if their consequence included coding or splice donor/acceptor mutations, max allele frequency was <0.01 in each of the large sequencing studies (ExAC, GnomAD/exomes, 1000 Genomes, ALFA) and predicted to be deleterious by both CAROL and FATHHM prediction tools (Lopes et al., 2012; Shihab et al., 2013). Variants called by targeted panel sequencing were called at a mean read depth of 278 (Standard Error of Mean =11). Exome studies were performed at mean depth of 40x. Pathogenic variants required a variant allele frequency ≥10%, a minimum read depth ≥10 and a minimum 2 variant forward reads and 2 variant reverse reads. Variants were further curated for obvious artefacts by visual inspection in Integrative Genomics Viewer (IGV) (J. T. Robinson et al., 2011). Chromosome-arm level copy-number estimates where derived from DNA methylation array data using conumee (R/Bioconductor). A larger previously published MB_Grp3_/MB_Grp4_ cohort (Sharma et al., 2019) (Schwalbe et al., 2017) (GSE130051 & GSE93646) to which 166 novel profiles were added (E-MTAB-10754) (n=1670, exact samples used are detailed in Supplementary Table 2) was used for methylation-only analysis.

#### RNA-seq analysis

Total RNA was extracted from snap frozen tissue samples using trizol extraction followed by Qiagen RNeasy Cleanup Kit and then subjected to transcriptome sequencing using Illumina TruSeq RNA Library Prep and HiSeq 2500 platform achieving a ∼90M paired end reads per sample. Following QC checks (fastqc/bamqc) samples were aligned to genome hg19 using *RNA-star* (Dobin et al., 2013) in two-pass alignment mode and per gene read counts generated using *ht-seq count* (Anders et al., 2015) and Gencode v25. Where isoform abundance estimates were required these were generated using *kallisto* (Bray et al., 2016). For differential expression analysis *DESeq2* (Love et al., 2014) (R/Bioconductor) was used for other analysis, clustering and visualization. Read counts were first normalized and a variance stabilizing transform was first applied using the *vst* function within *DESeq2* (R/Bioconductor). Additionally, a batch correction controlling for sequencing batch was applied using the implementation of ComBat within the *sva* package (R/Bioconductor). Consensus NMF analysis was performed as per the method described in Schwalbe *et al* (Schwalbe et al., 2017) and Sharma *et al* (Sharma et al., 2019). Briefly, multi-run NMF is performed with n=250 iterations of 80% bootstrapping. Metagenes calculated following each iteration are projected on to each removed sample and k-means clustering used to predict the class of each removed sample based on the larger training set. A range of NMF metagene ranks (3-10) and k-means clusters (3-10) are tested and cophenetic indices (a shorthand measure of the robustness of sample clustering) used to evaluate the consistency of classification for each combination of metagenes. Samples which were assigned to the same class with <90% consistency upon resampling were designated as MB-NOS, except where they were alternately assigned as MB_Grp3_ or MB_Grp4_ with >90% consistency, in which case they were classified as MB_Grp3_/MB_Grp4_.

Averaged and standardized metagene h-values from across the bootstraps were used as measures of metagene expression. All NMF projections were performed using column-rank and post-projection normalization as per the method described by Tamayo *et al* (Tamayo et al., 2007). t-SNE were used for visualization was performed using the *Rtsne* package (R/CRAN).

G3/G4 score was calculated by applying a logistic transformation 1/(1+exp(-x)) to the MB_Grp3_ and MB_Grp4_ metagenes (excluding two outliers). The G3/G4 score was calculated as the MB_Grp3_ proportion of the total metagene scaled to between 0 and 1. For convenience of visualization, or where categorical comparison was required, we referred to individuals >0 & ≤0.2 as “HighG4”, >0.2 & ≤0.4 as “LowG4”, >0.4 & ≤0.6 as “G3.5”, >0.6 & ≤0.8 as “LowG3” and >0.8 & ≤1 as “HighG3”.

RNA editing was estimated using the QEdit/Reditools pipeline as previously described (https://github.com/BioinfoUNIBA/QEdit) (Giudice et al., 2020). Differential RNA-editing was calculated using a p-adjusted (Benjamini-Hochberg) Mann-Whitney U-test for two group analysis and Anova with TukeyHSD (post-hoc) for multi-group analysis. Where unknown from DNA analysis *GFI1/GFI1B*, *PRDM6* rearrangements were each inferred from RNA-seq data as per the method used originally by Northcott *et al* (Northcott et al., 2017; 2014).

GSEA was performed using MsigDb library version 7.1 and the implementation of the original algorithm within the package *fgsea* (R/Bioconductor) and ssGSEA using the implementation within *GSVA* (R/Bioconductor) (Hänzelmann et al., 2013). The following gene sets were selected as reflective of the pathway categories given in Figure 4C. MYC = “HALLMARK_MYC_TARGETS_V2”, “MYC_UP.V1_UP”, “DANG_MYC_TARGETS_UP”. Cell Cycle = “FISCHER_G1_S_CELL_CYCLE”, “GO_POSITIVE_REGULATION_OF_CELL_CYCLE”, “GO_SIG NAL_TRANSDUCTION_INVOLVED_IN_CELL_CYCLE_CHECKPOINT”, TP53 = “CEBALL OS_TARGETS_OF_TP53_AND_MYC_UP”, “REACTOME_TRANSCRIPTIONAL_REGULATION_BY_TP53”, “REACTOME_TP53_REGULATES”, MTOR = “HALLMARK_MTORC1_SIGNALING”, “MTOR_UP.V1_UP”, “MTOR_UP.N4.V1_UP”, PHOTORECEPTOR = “GO_EYE_PHOTOREC EPTOR_CELL_DIFFERENTIATION”, “GO_CAMERA_TYPE_EYE_PHOTORECEPTOR_CELL_ DIFFERENTIATION”, “GO_EYE_PHOTORECEPTOR_CELL_DEVELOPMENT”, TGFB1 = “KARL SSON_TGFB1_TARGETS_UP”, “JAZAG_TGFB1_SIGNALING_VIA_SMAD4_UP”, “KARAKAS_ TGFB1_SIGNALING” NOTCH = “GO_POSITIVE_REGULATION_OF_NOTCH_SIGNALING_PATHWAY”, “REACTOME_ACTIVATED_NOTCH1_TRANSMITS_SIGNAL_TO_THE_NUCLEUS”, “NGUYEN_NOTCH1_TARGETS_UP”, Neuronal Diff = “GO_CENTRAL_NERVOUS_SYSTEM_NEURON_DIFFERENTIATION” “LE_NEURONAL_DIFFERENTIATION_UP”.

In analyzing association with G3/G4 score, the loss or gain of each non-acrocentric chromosome arm was considered as were the more frequent MB_Grp3_/MB_Grp4_ mutations in genes *ATM*, *CTDNEP1*, *KDM6A*, *KIF26B*, *KMT2C*, *KMT2D*, *NBAS*, *NEB*, *RYR3*, *SMARCA4*, *SPTB*, *TBR1*, *TSC2*, and *ZMYM3*.

#### DNA methylation analysis

Beta/M-values were derived from HumanMethylation450 BeadChip (450k) and Infinium HumanMethylationEPIC (850k) arrays using the ssNOOB method within the package *minfi* (Aryee et al., 2014) excluding known SNPs and cross-hybridizing probes. In order to construct a random forest classifier which predicted G3/G4 score from DNA methylation data, we performed feature selection of CpGs using 192 MB_Grp3_/MB_Grp4_ samples with both RNA-seq (i.e. known G3/G4 score) and Methylation array. We constructed using *limma* (R/Bioconductor) a number of bootstrapped (80% with 100 iterations) significance tests testing differential methylation between each of the categories HighG4, LowG4, G3.5, LowG3 and HighG3. We measured average performance for a range of numbers of features (10-100) on removed samples using a tuned support vector machine, however performance plateaued after a certain number of features, so it was decided to select the top 80 most frequently selected CpGs for each comparison. Thus n=400 CpG features were used to train a random forest classifier which was then subject to recursive feature elimination using 50x cross-validation and implemented using the *rfe/rfeControl* function within the *caret* package (R/CRAN). Where sigmoid curves are shown, these were fitted using the *fitmod* function within the *DoseFinding* package (R/Bioconductor). For visualization these were scaled to a minimum 0 and maximum 1.

Methylation subtype calling (Sharma et al., 2019) was obtained using an extension of the Heidelberg brain tumor classifier available at [https://www.molecularneuropathology.org/mnp]. A methylation classifier prediction score of >0.8 was used to assign subtype. Samples were excluded if not confirmed as MB by MNP. Significantly differentially methylated regions (DMRs) distinguishing G4High, G4Low, G3.5, G3Low and G3High were calculated using *dmrcate* (R/Bioconductor) using settings lambda=1000, C=2. Regions were considered when the total number of CpGs ≥ 5, the minimum FDR < 0.05 and the mean Beta fold change > 0.25. These were further filtered to identify DMRs which overlapped with the MB_Grp3_/MB_Grp4_ specific enhancer/superenhancer regions identified by Lin *et al* (Lin et al., 2016).

#### scRNA-seq analysis

Previously published medulloblastoma scRNA-seq dataset (Hovestadt et al., 2019) GSE119926 was used. However, we used only the MB_Grp3_/MB_Grp4_ primary patient samples (excluding the patient-derived xenografts) (n=4256 cells, n=15 samples) and excluded patients SJ970 and SJ723 due to the relatively few available cells. The pre-publication Human fetal cerebellar single cell reference data set, consisting of 69,174 cells, classified into 21 cell types and derived from 15 donors between 9 and 21 PCW, details can be found within https://www.biorxiv.org/content/10.1101/2020.06.30.174391v1 (Aldinger *et al* in press Nature Neuroscience). For the purposes of metagene projection *Seurat* (R/Bioconductor) (Butler et al., 2018) was used to select the 5000 most variable features using the “vst” method for both data sets and the resulting normalized matrices subject to NMF projection of the bulk metagenes and calculation of the G3/G4 score as per the bulk analysis described above. In this way, a per-cell metagene score and G3/G4 score was calculated.

Developmental trajectory analysis was performed using *monocle v3* (Qiu et al., 2017) (R/Bioconductor) using 12,243 cells classified as RL, GCP, GN or eCN/UBC which we defined broadly as the rhombic lip lineage as per Aldinger *et al*. Monocle v3 functions used were preprocess_cds, align_cds, reduce_dimension, cluster_cells, learn_graph, order_cells and plot_cells to visualize by UMAP. The relevant branches for the MB_Grp3_/MB_Grp4_ and MB_SHH_ were divided as indicated (Figure 8) and the relationship between pseudotime and G3/G4 score/metagene was defined using a loess curve function. This enabled developmental and oncogenic events to be mapped onto a common scale (Figure 8). Genes whose expression varied significantly according to pseudotime were detected using Moran’s test statistic as implemented by *monocle v3*. For analysis of the differences between MB_SHH-Infant_ and MB_SHH-Child,_ a further metagene calculated using NMF rank = 2 only on MB_SHH_ (67/331 samples) was additionally projected onto the single cells in the same manner as the other metagenes. For calculating empirical density, the *density* function was used (R/Bioconductor) except where weighted two-dimensional estimation was needed in which case the *kde2d.weighted* function from the package *ggtern* (R/Bioconductor) was used. Weights were calculated as the number of cells at a given sampling point (9-21PCW) as a proportion of the total number of cells sampled.

### Quantification and Statistical Analysis

Data analysis and visualization was carried out in R 3.5.3 except for the analysis of fetal cerebellar scRNA-seq which was performed using R 4.0.2. CRAN and Bioconductor packages used are given in the key resources table. To test significant association with time to death/progression free survival a log-rank test or cox-regression was used. Association analysis of individual clinico-pathological mutational and copy number features with G3/G4 score was performed using Kolmogorov-Smirnoff test. Where gene expression/pathway associations with G3/G4 score are assessed, these are assessed using Pearson’s correlation coefficient. The test statistics and significant P-values (p<0.05) are stated in text and figures and were adjusted for multiple hypothesis testing using Benjamini-Hochberg where appropriate throughout. Where values of n are given, these generally pertain to number of sample/individual patients except where otherwise indicated. Boxplots, where used, show dispersion as per standard i.e. (center line = median, box = interquartile range, whisker= range minus outliers).

Data were excluded where samples were clearly indicated to be duplicated across multiple related datasets. Additional exclusions were carried out for samples where methylation array detection P value did not reach significance threshold in at least 90% of the array. Methylation samples were excluded from the analysis if not confirmed as medulloblastoma by MNP2.0. In our analysis of the scRNA-seq dataset GSE119926 we excluded patients SJ970 and SJ723 due to the relatively few available cells.

### Key Resources Table

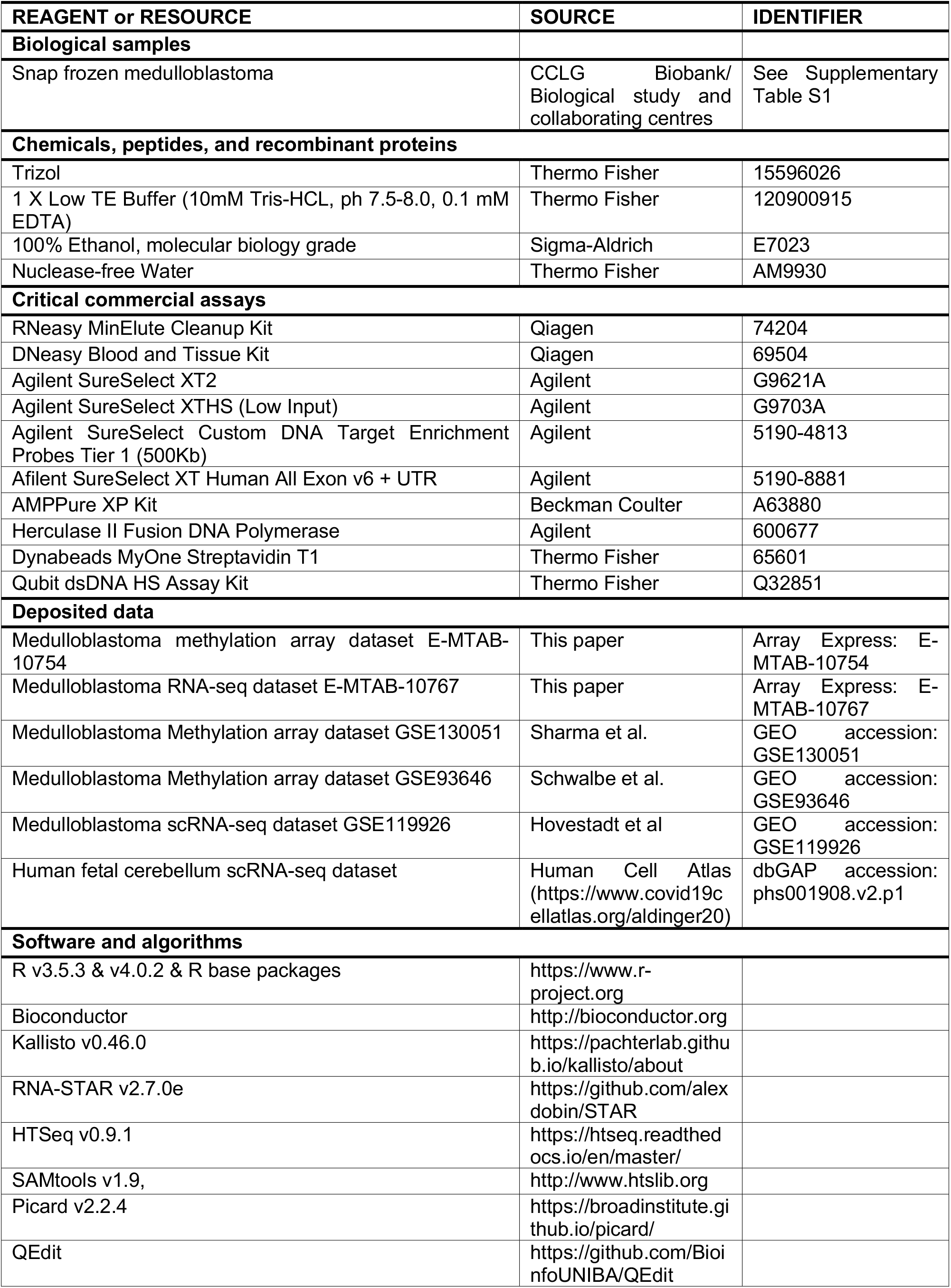
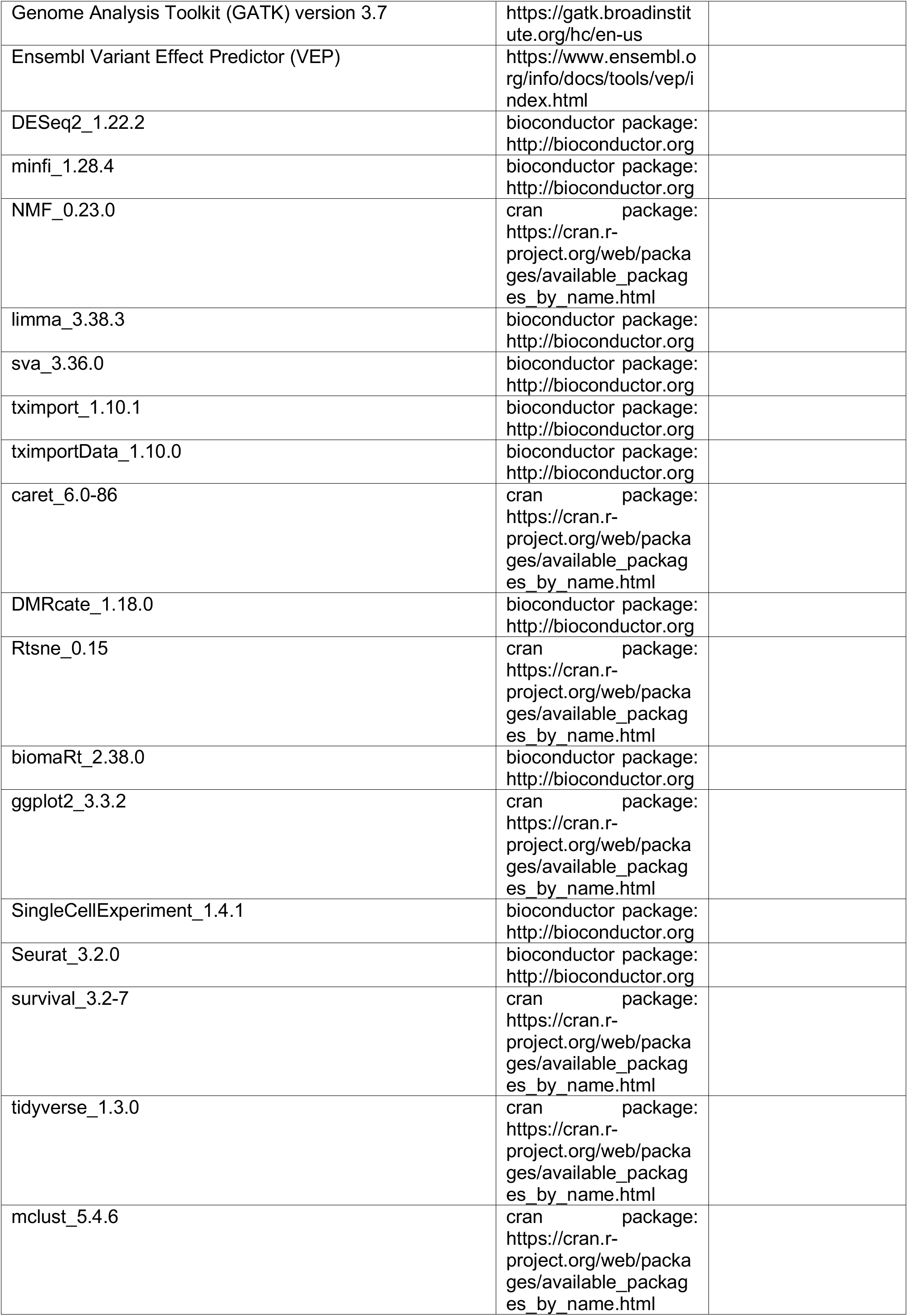
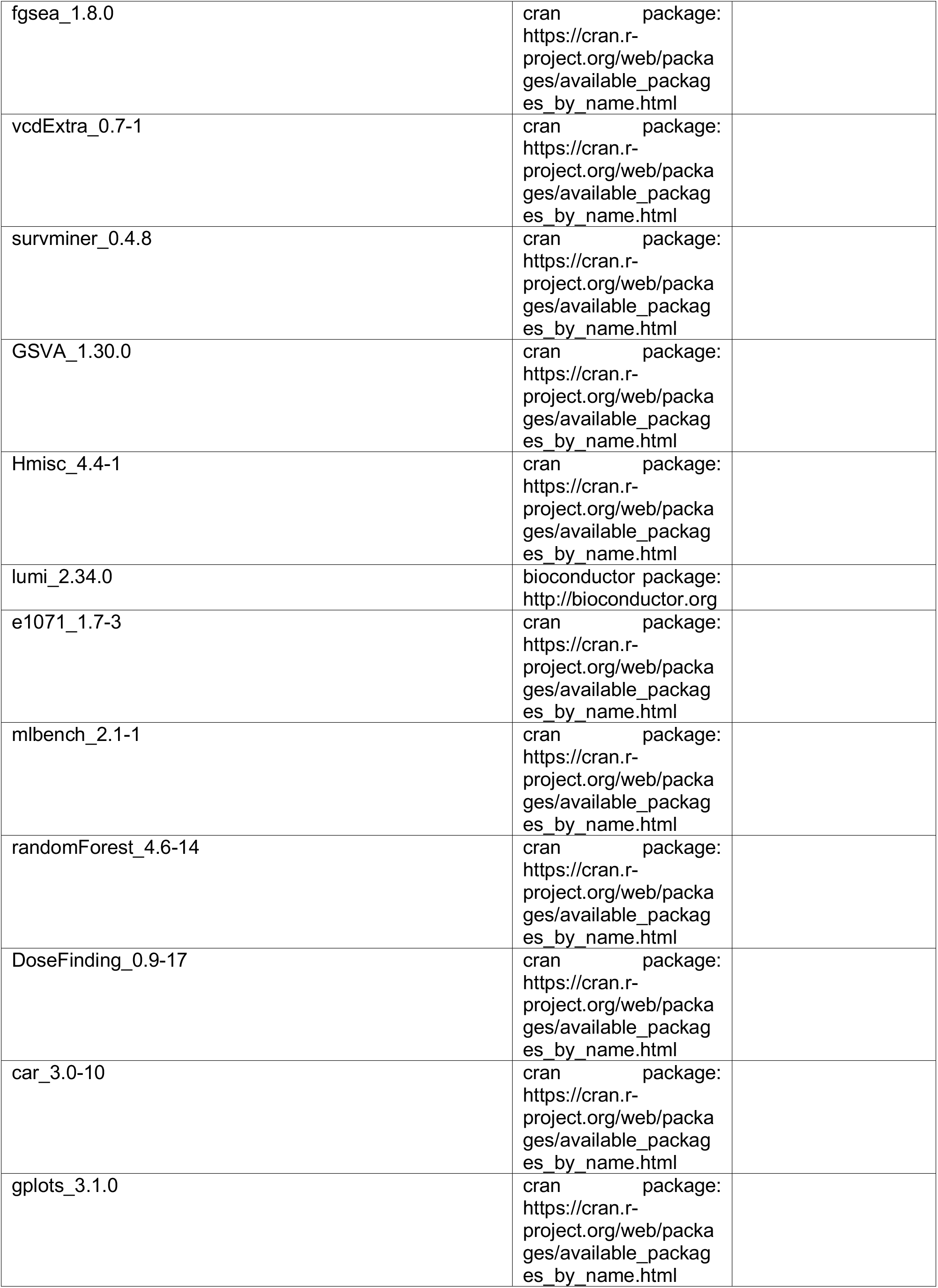
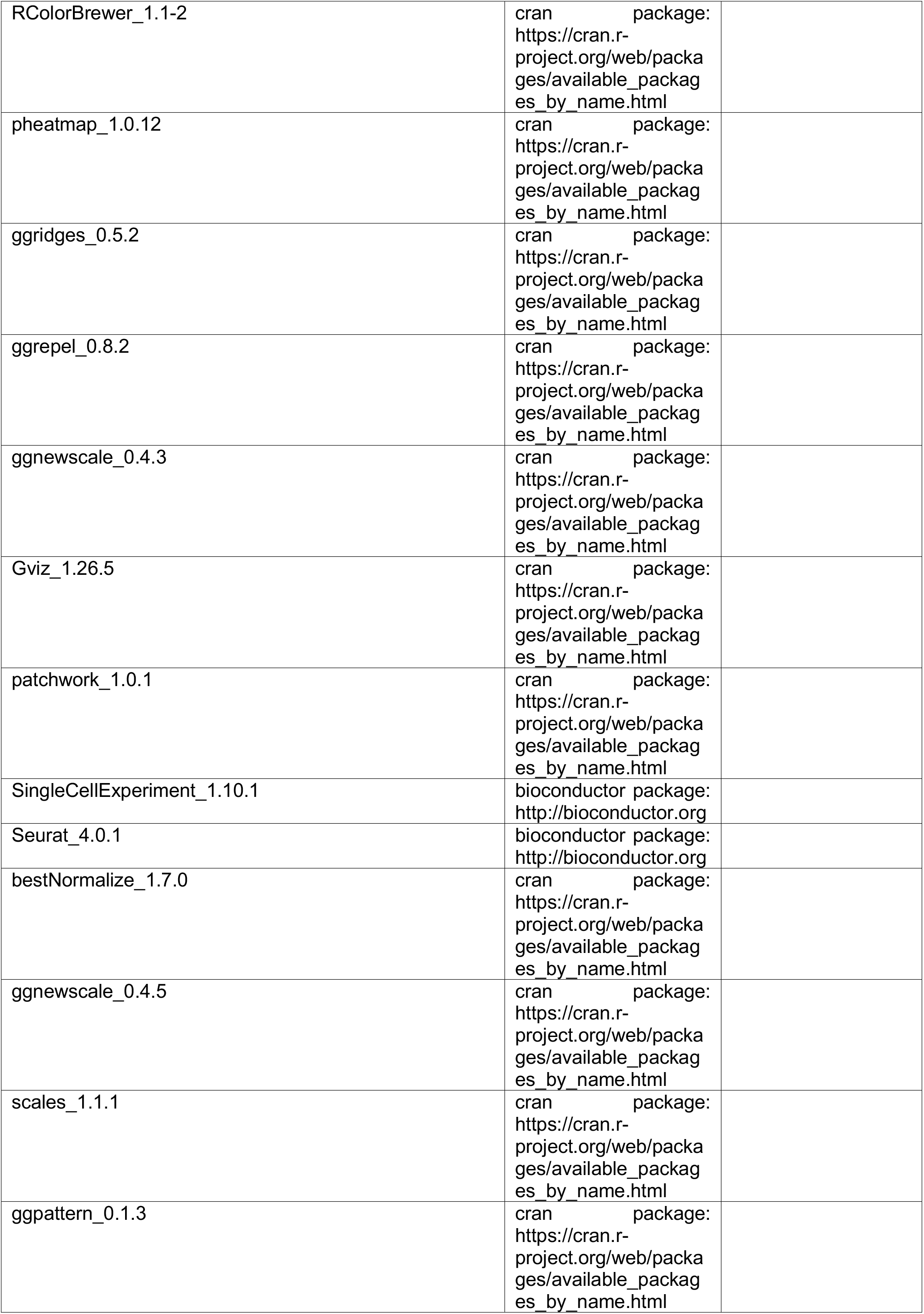
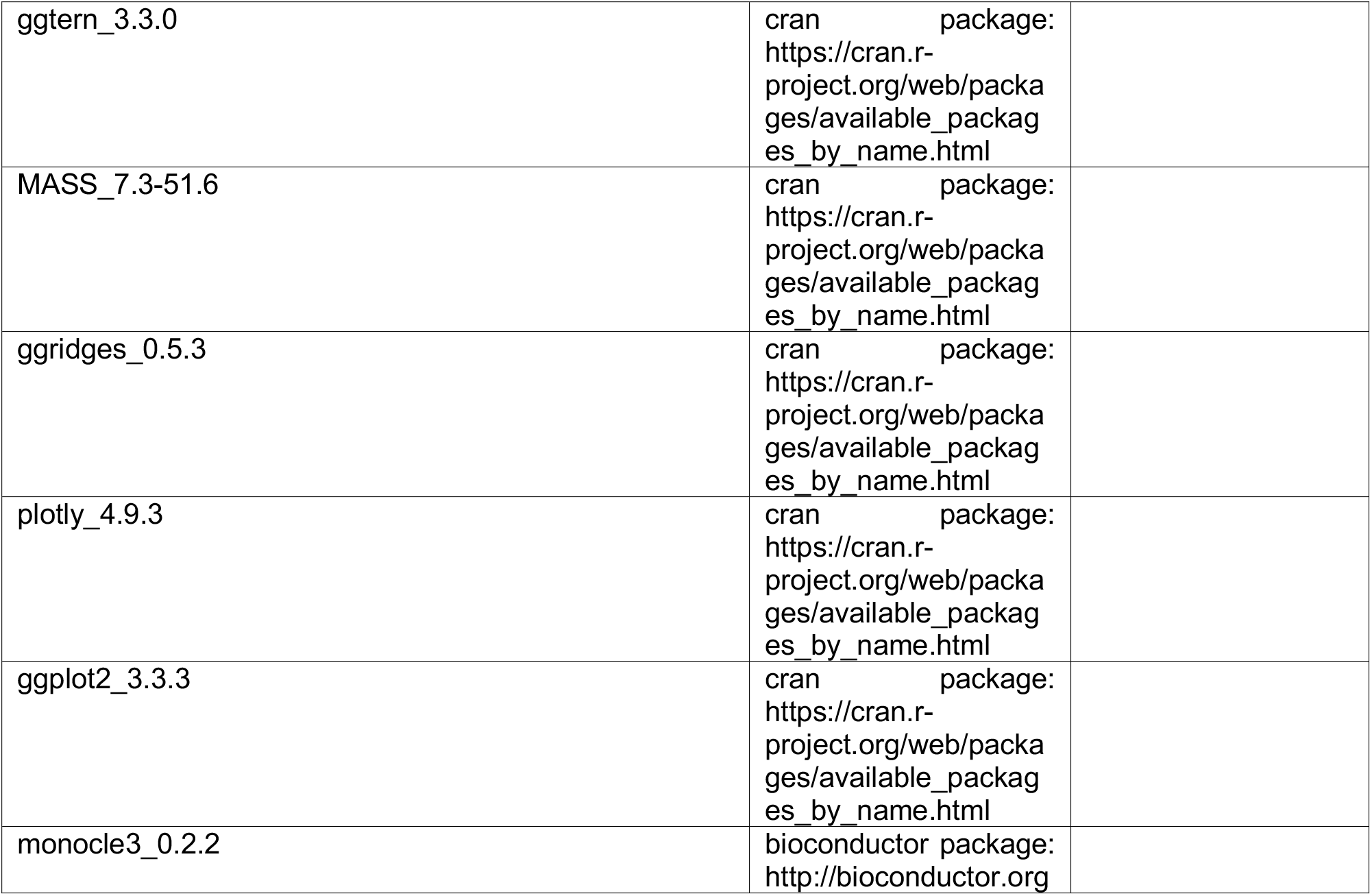

## Supplemental information

Supplementary Table 1. Demographic summary of patients.

Supplementary Table 2. Sentrix IDs of samples used in larger methylation cohort.

**Figure 1S.**
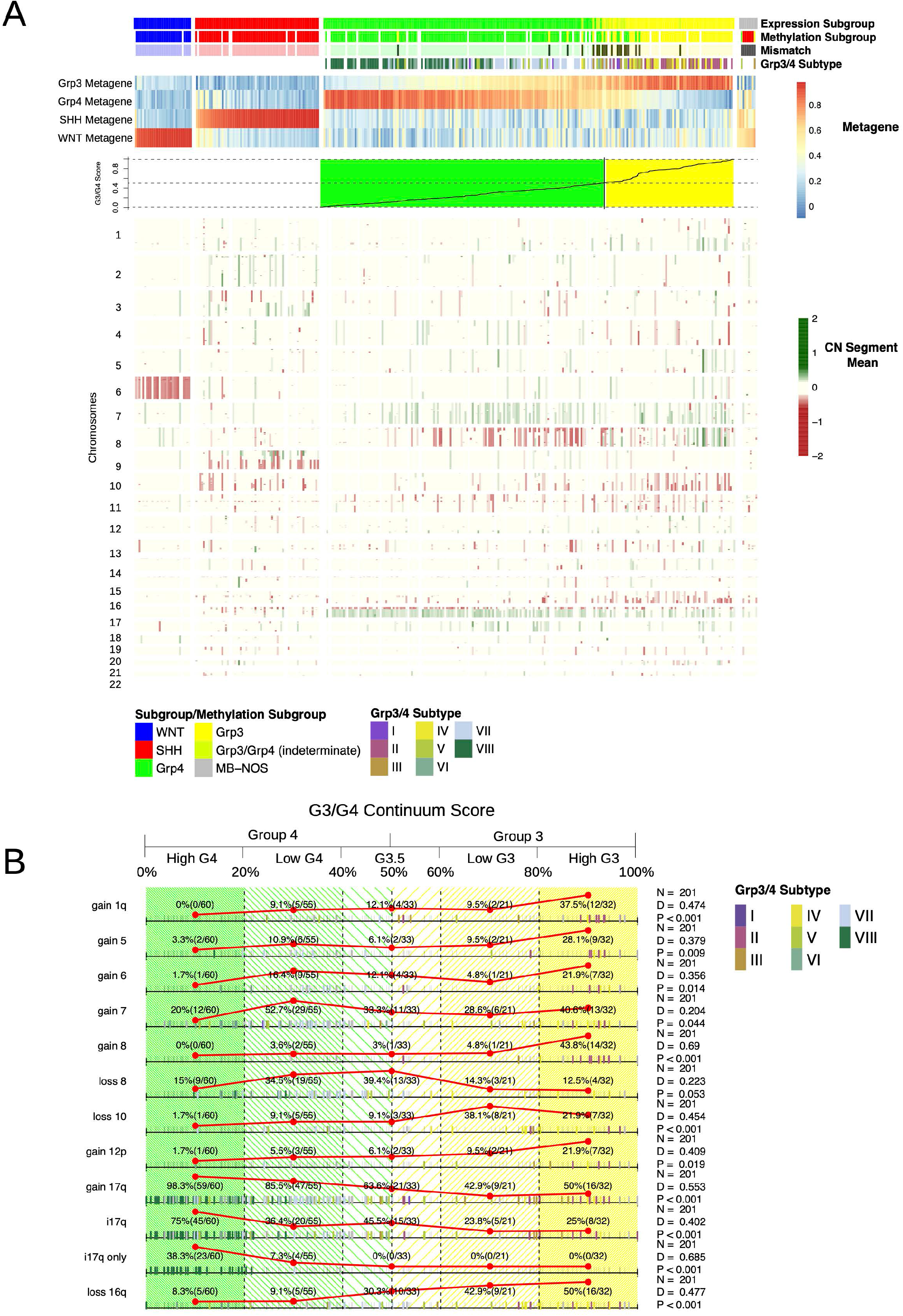
**A:** Heatmap showing copy number changes by chromosome for n=331 MB and grouped by subgroup. MB_Grp3_/MB_Grp4_ individuals are ordered by G3/G4 score. Column annotation shows subgroup as determined by RNA-seq (Expression Subgroup) as determined by DNA methylation array (Methylation Subgroup), and DNA methylation MB_Grp3_/MB_Grp4_ subtype (I-VIII) as per Sharma *et al* 2019(Sharma et al., 2019) as defined using MNPv2 classifier(Capper et al., 2018) (Grp3/4 Subtype). The line plot (bottom) shows G3/G4 score. **B:** Rug plot showing distribution of aneuploidy/copy number change with respect to G3/G4 score. Summary counts are given according to the convenient divisions of HighG4, LowG4, G3.5, LowG3, HighG3 and reflected by the red line plots. Presence of a given feature is indicated by a bold tick mark, the color of which indicates methylation MB_Grp3_/MB_Grp4_ subtype (I-VIII). Adjusted P-values for a Kolmogorov-Smirnoff statistic (D) are shown to denote non-random distribution of features with respect to G3/G4 score.

**Figure 2S.**
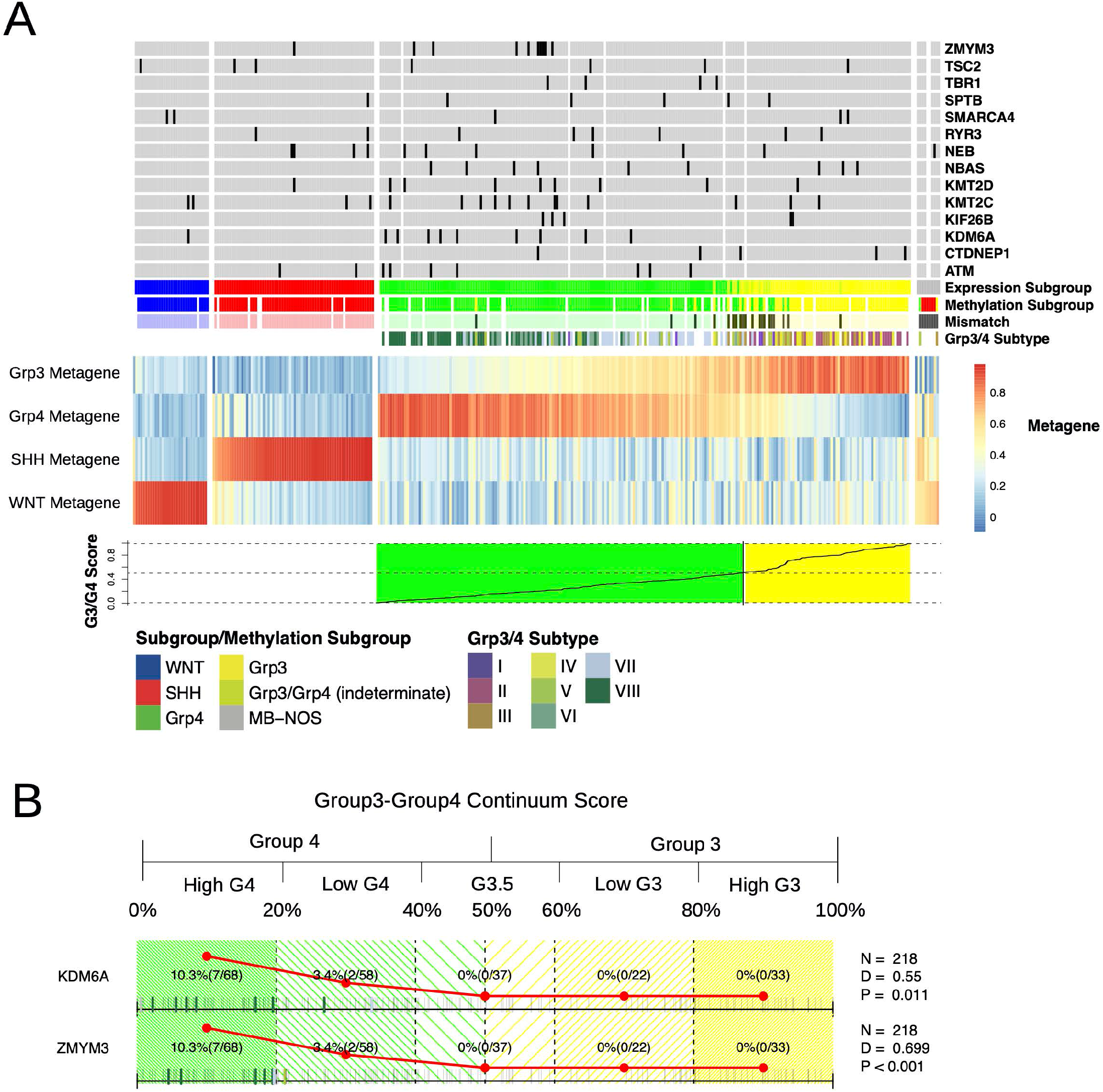
**A:** Heatmap showing 4 consensus NMF metagenes calculated for n=331 MB and grouped by subgroup. MB_Grp3_/MB_Grp4_ individuals are ordered by G3/G4 score. Column annotation shows subgroup as determined by RNA-seq (Expression Subgroup) as determined by methylation (Methylation Subgroup), methylation MB_Grp3_/MB_Grp4_ subtype (I-VIII) as per Sharma *et al* 2019 as defined using MNPv2 classifier (Grp3/4 Subtype). Presence of mutations are indicated to be present or not by dark grey shading. White indicates missing data. **B:** Rug plot showing distribution of mutations with respect to G3/G4 score. Summary counts are given according to the convenient divisions of HighG4, LowG4, G3.5, LowG3, HighG3 and reflected by the red line plots. Presence of a given feature is indicated by a bold tick mark the color of which indicates methylation MB_Grp3_/MB_Grp4_ subtype (I-VIII). P-values for a Kolmogorov-Smirnoff statistic (D) are shown to denote non-random distribution of features with respect to G3/G4 score.

**Figure 3S.**
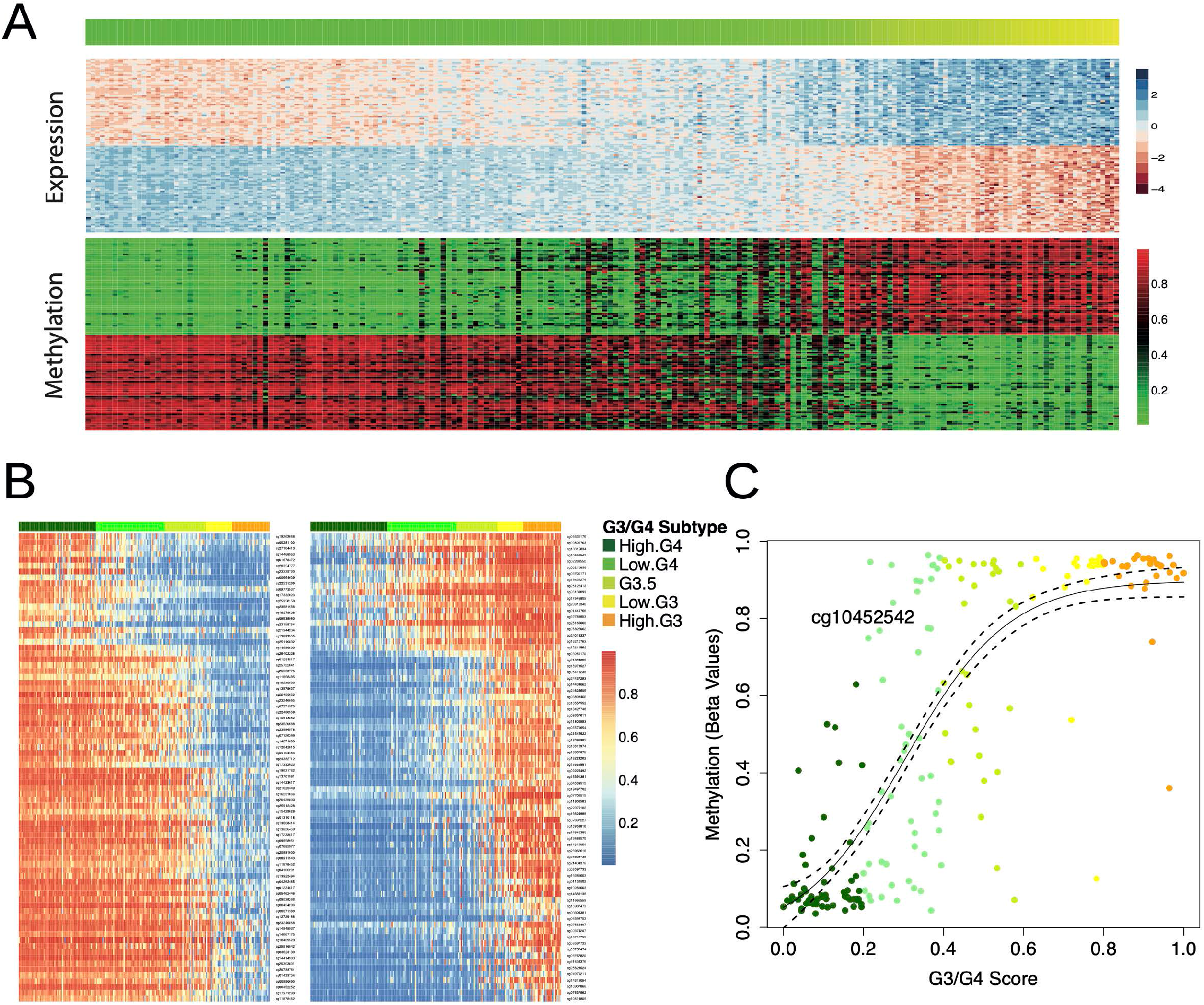
**A:** Heatmap showing top 50 genes most significantly differentially expressed (top) and top 50 CpGs differentially methylated between MB_Grp3_ and MB_Grp4_. Samples are ordered according to G3/G4 score. Note the difference in gradation for the expression values as opposed to the more binary distribution of DNA methylation beta-values. **B:** Heatmap showing DNA methylation values of the top 40 most discriminatory CpGs distinguishing HighG4 (dark green), LowG4 (light green), Low G3 (yellow) and High G3 (orange). G4 hypermethylated CpGs are shown on the left and hypomethylated CpGs on the right. Samples are ordered according to G3/G4 score and G3/G4 categories (HighG4, LowG4, G3.5, LowG3, HighG3) are annotated. **C:** Scatterplot showing beta-values for CpG “cg19784198” colored by G3/G4 categories (HighG4, LowG4, G3.5, LowG3, HighG3) an example of a CpG differentially expressed between MB_Grp3_ and MB_Grp4_ showing a bimodal methylation distribution. The relationship with G3/G4 score can effectively be modelled by a sigmoid/logistic function.

**Figure 4S.**
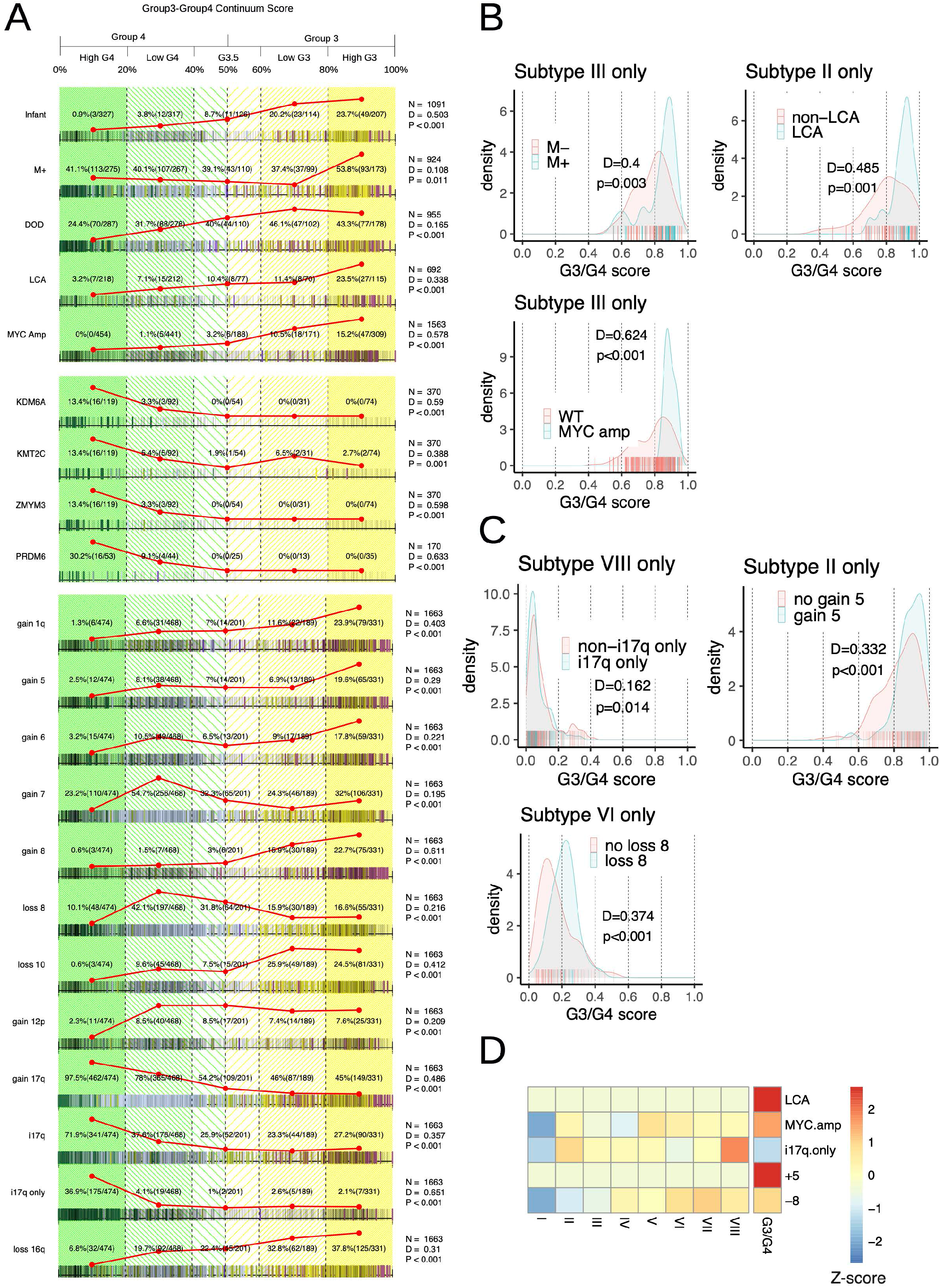
**A:** Rug plot showing distribution of clinicopath features (top) mutations (middle) and copy number (bottom) with respect to G3/G4 score derived from DNA methylation data. Summary counts are given according to the convenient divisions of HighG4, LowG4, G3.5, LowG3, HighG3 and reflected by the red line plots. Presence of a given feature is indicated by a bold tick mark the color of which indicates methylation MB_Grp3_/MB_Grp4_ subtype (I-VIII). P-values for a Kolmogorov-Smirnoff statistic (D) are shown to denote non-random distribution of features with respect to G3/G4 score. Infant=age at diagnosis < 3 years, Metastases = M+, DOD=Dead of Disease, LCA = Large Cell Anaplasia, PRDM6 = *PRDM6* rearrangement. **B:** Empirical density and rug plots showing the distribution of M+ in MB_Grp3_/MB_Grp4_ subtype III, LCA in MB_Grp3_/MB_Grp4_ subtype II and *MYC* amplification in MB_Grp3_/MB_Grp4_ subtype III with respect to G3/G4 score. The given clinico-pathological features are significantly non randomly distributed with respect to G3/G4 score even within specific MB_Grp3_/MB_Grp4_ subtypes as shown by Kolmogorov-Smirnoff test (D). **C:** Empirical density and rug plots showing the distribution of copy number changes i17q in MB_Grp3_/MB_Grp4_ subtype VIII, Gain of chromosome 5 in MB_Grp3_/MB_Grp4_ subtype II and loss of chromosome 8 in MB_Grp3_/MB_Grp4_ subtype VI with respect to G3/G4 score. The given copy number features are significantly non randomly distributed with respect to G3/G4 score even within specific MB_Grp3_/MB_Grp4_ subtypes as shown by Kolmogorov-Smirnoff test (D). **D:** Heatmap showing (z-scores) of either methylation MB_Grp3_/MB_Grp4_ subtypes (I-VIII) or G3/G4 score as an explanatory variable in a logistic regression determining presence or absence of the indicated clinico-path, copy number changes or mutational characteristics.

**Figure 5S.**
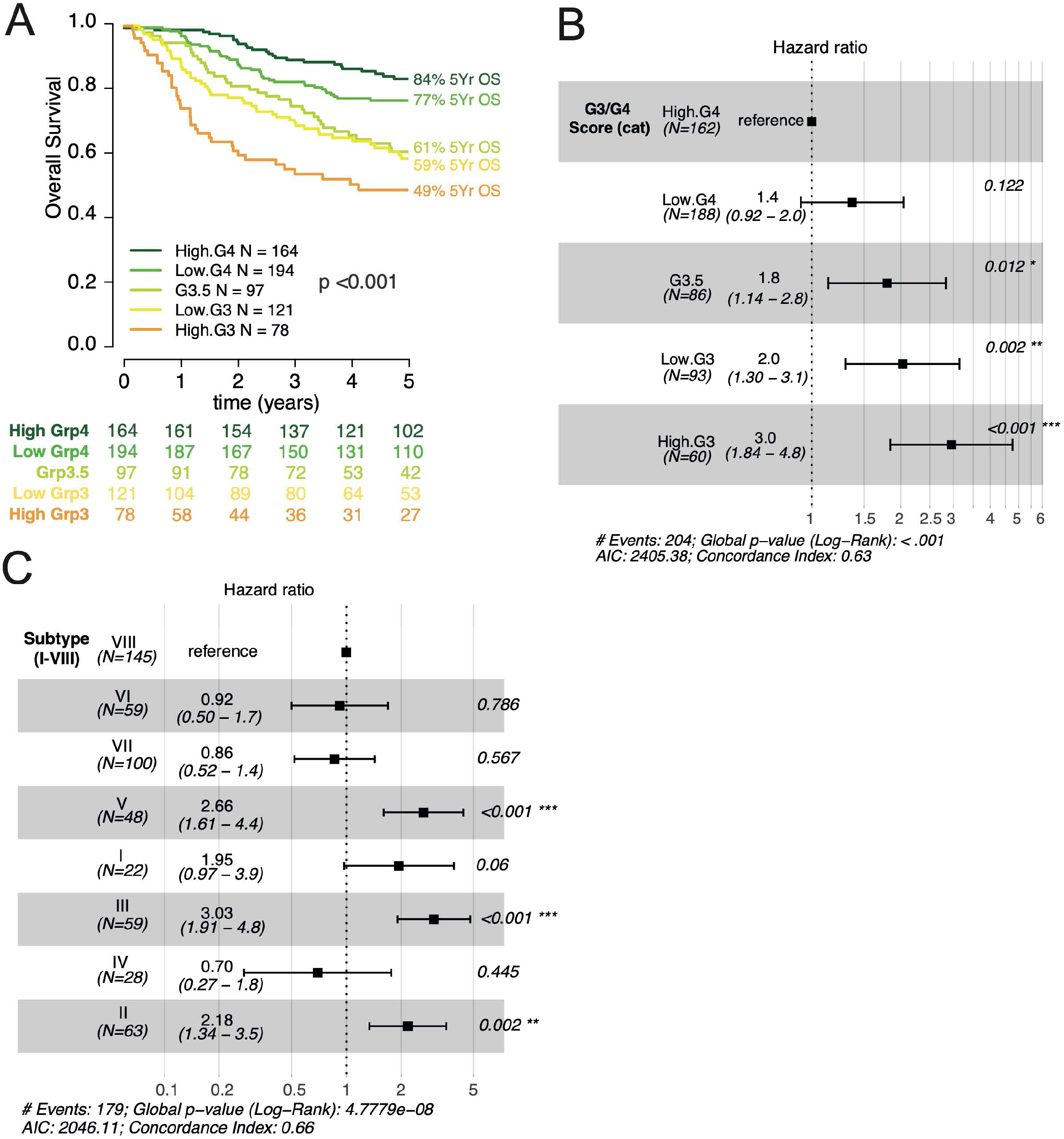
**A:** Kaplan-Meier plot showing significant differences in MB_Grp3_/MB_Grp4_ overall survival (patients of all ages) by G3/G4 continuum position. **B:** Forest plot showing univariate Cox models (patients > 3 years) of overall survival containing the variables G3/G4 score (as predicted by DNA methylation) treated as a categorical variable and **C:** MB_Grp3_/MB_Grp4_ methylation subtype.

**Figure 6S.**
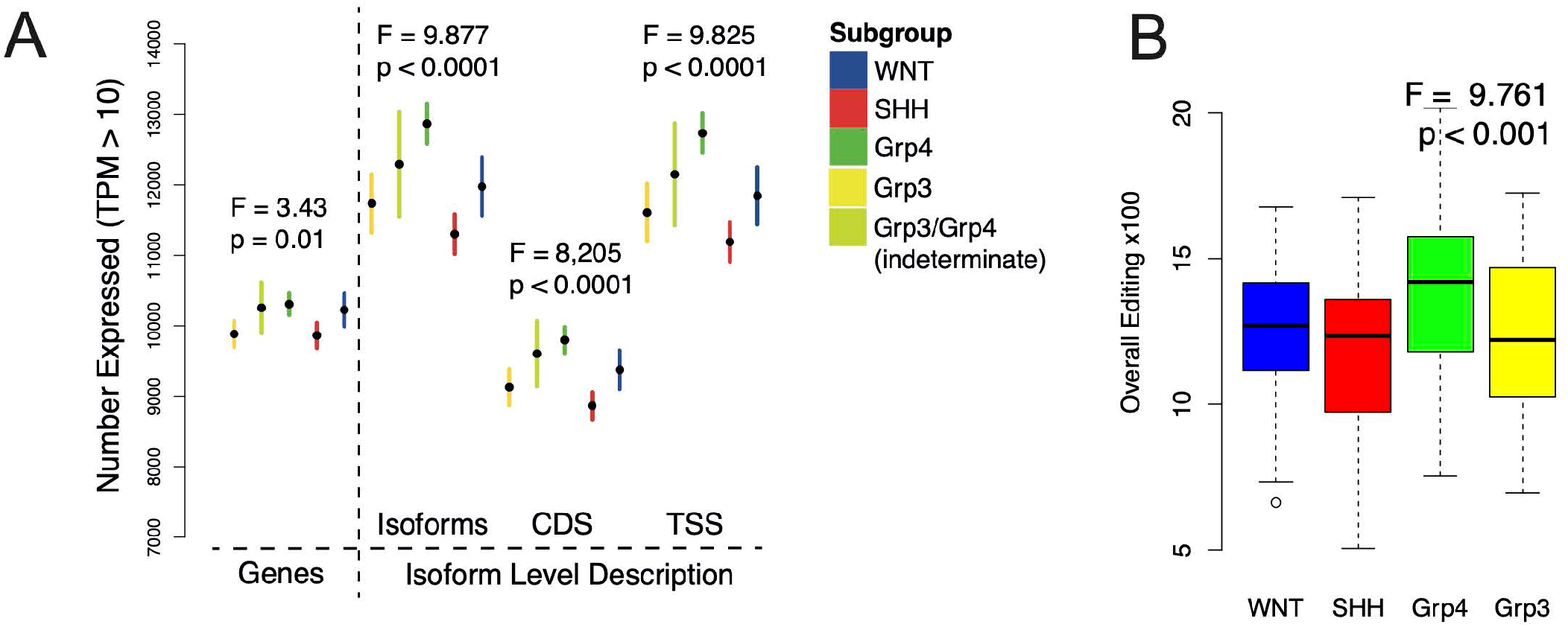
**A:** Boxplot showing the distribution by MB subgroup of moderately expressed genes, isoforms, CDS or TSS as defined by a TPM>10. **B:** Boxplot showing significant differences in OEI (Overall Editing Index), i.e. level of RNA-editing by MB subgroup.

**Figure 7S.**
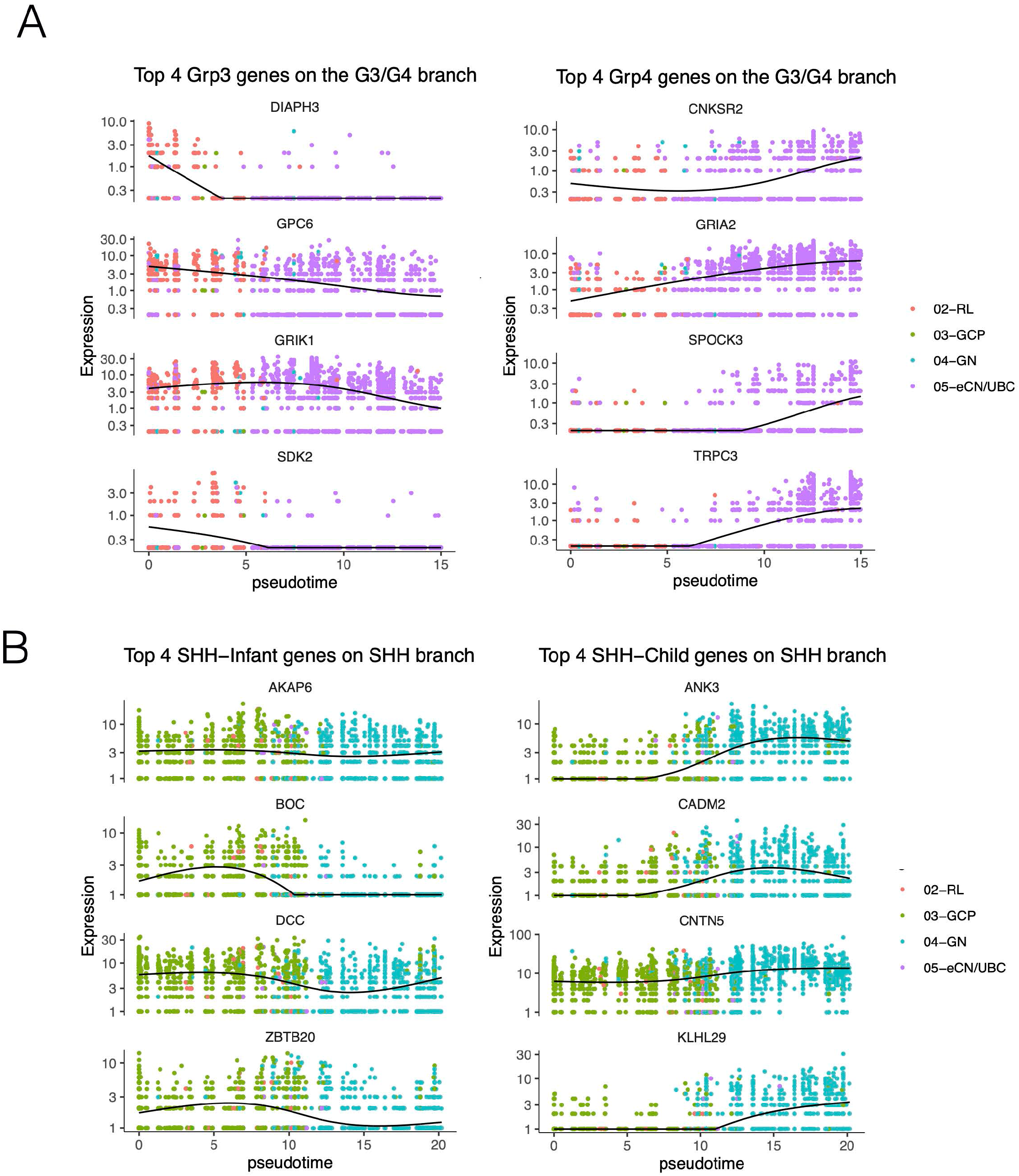
Plots showing the per-cell expression of genes whose expression varies according to pseudotime on the **A:** RL to eCN/UBC branch (MB_Grp3_ specific genes are shown on the left and MB_Grp4_ specific genes shown on the right) and the **B:** GCP to GN branch (MB_SHH-Infant_ specific genes are show on the left and MB_SHH-Child_ specific genes are shown on the right). Cell type is denoted by color. Black line represents a loess curve. Expression is represented as normalized count data.

**Figure 8S.**
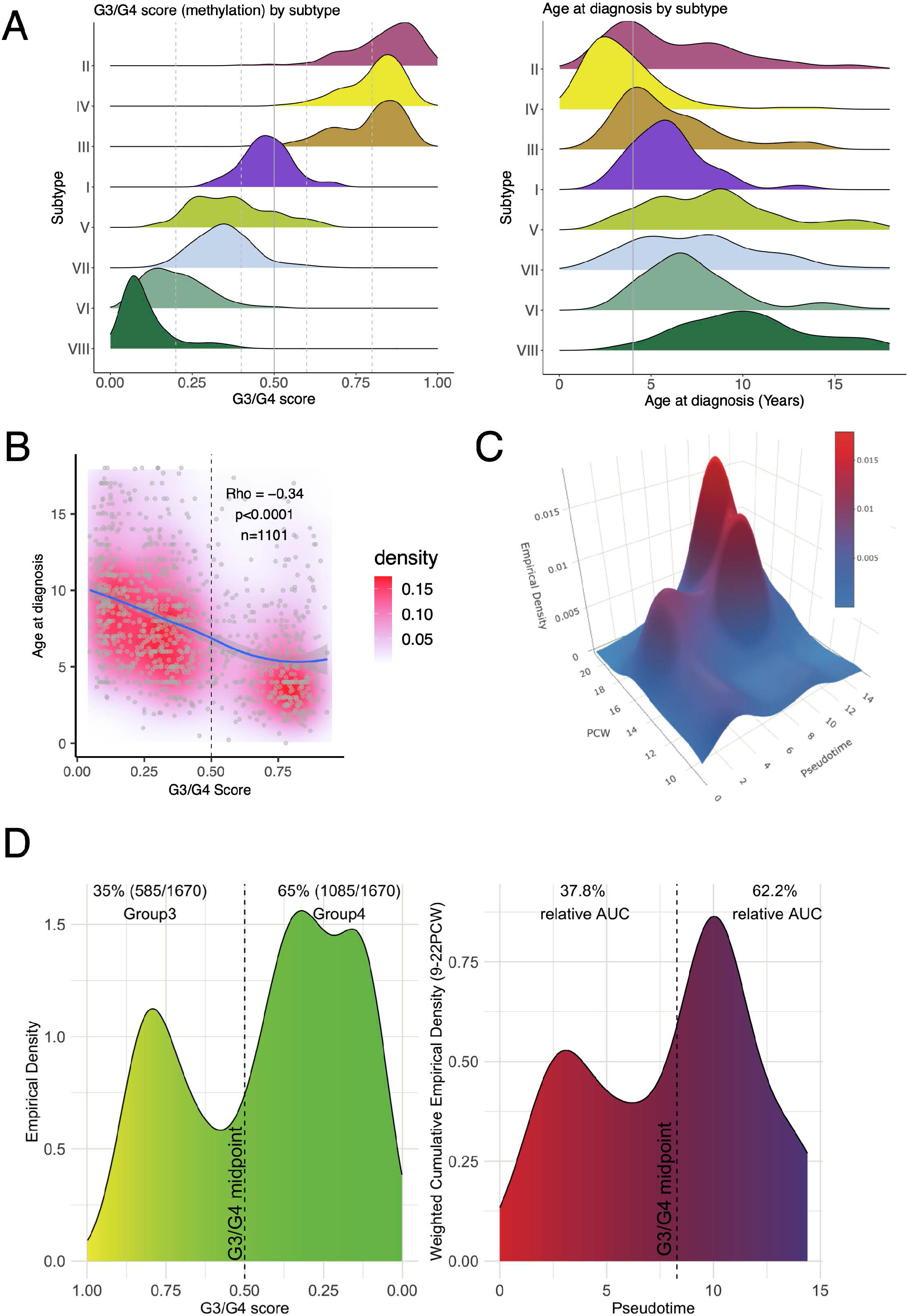
**A:** Ridgeplots showing (left) distribution of G3/G4 score MB_Grp3_/MB_Grp4_ patients by methylation subtype (I-VIII) (right) and distribution of age at diagnosis by DNA methylation subtype (I-VIII). **B:** Scatterplot showing age at diagnosis by G3/G4 score (as determined by DNA methylation), 2d empirical density is shown as red shading and a loess curve with 95% CI is shown as blue line with grey shading. **C:** Plot showing 3d weighted empirical density of cells on the RL to eCN/UBC branch in fetal cerebellar development from 9-21 PCW and by pseudotime. Cells were weighted according to the proportion of cerebellar cells sampled at that time point. **D:** Empirical density plots showing (left) distribution of MB_Grp3_/MB_Grp4_ patients by G3/G4 score and (right) the empirical density cumulatively across 9-21 PCW (essentially the 3d volume under the plane shown in panel C) by pseudotime. Note the similarities in distribution. The dotted line shows the divide between Grp3 and Grp4 space on both G3/G4 and pseudotime scales and the percentages relate to the relative amounts of density represented on either side of the divide.

