## Supplementary Table 1 for "Medulloblastoma Group 3 and 4 Tumors Comprise a Clinically and Biologically Significant Expression Continuum Reflecting Human Cerebellar Development"

### Demographic Summary of Patients

| | Subgroup | | | | | | |
|---|---|---|---|---|---|---|---|
| | Group3 | Group3/Group4 | Group4 | SHH | WNT | MB-NOS | Total |
| **Sex** | | | | | | | |
| Male | 41/55 (74.5%) | 8/12 (66.7%) | 96/132 (72.7%) | 31/61 (50.8%) | 13/28 (46.4%) | 5/7 (71.4%) | 194/295 (65.8%) |
| **Age** | | | | | | | |
| Adult (>16 years) | 0/56 (0.0%) | 0/12 (0.0%) | 2/134 (1.5%) | 5/61 (8.2%) | 1/29 (3.4%) | 1/7 (14.3%) | 9/299 (3.0%) |
| Infant (<3 years) | 18/56 (32.1%) | 0/12 (0.0%) | 6/134 (4.5%) | 18/61 (29.5%) | 0/29 (0.0%) | 2/7 (28.6%) | 44/299 (14.7%) |
| **Clinicopathology** | | | | | | | |
| Relapse | 25/50 (50.0%) | 3/10 (30.0%) | 43/124 (34.7%) | 23/55 (41.8%) | 3/28 (10.7%) | 4/7 (57.1%) | 101/274 (36.9%) |
| DOD | 26/54 (48.1%) | 3/12 (25.0%) | 39/131 (29.8%) | 22/60 (36.7%) | 4/29 (13.8%) | 4/7 (57.1%) | 98/293 (33.4%) |
| Sub Total Resection | 8/50 (16.0%) | 2/12 (16.7%) | 31/114 (27.2%) | 4/55 (7.3%) | 1/28 (3.6%) | 0/6 (0.0%) | 46/265 (17.4%) |
| LCA | 7/42 (16.7%) | 2/11 (18.2%) | 8/105 (7.6%) | 9/50 (18.0%) | 4/24 (16.7%) | 0/6 (0.0%) | 30/238 (12.6%) |
| DN | 1/42 (2.4%) | 0/11 (0.0%) | 6/105 (5.7%) | 16/50 (32.0%) | 0/24 (0.0%) | 0/6 (0.0%) | 23/238 (9.7%) |
| M+ | 21/50 (42.0%) | 4/12 (33.3%) | 33/122 (27.0%) | 12/58 (20.7%) | 2/29 (6.9%) | 1/6 (16.7%) | 73/277 (26.4%) |
| **Mutations** | | | | | | | |
| GFI1B rearrangement | 0/63 (0.0%) | 0/13 (0.0%) | 3/147 (2.0%) | 0/67 (0.0%) | 0/31 (0.0%) | 0/10 (0.0%) | 3/331 (0.9%) |
| GFI1 rearrangement | 5/63 (7.9%) | 0/13 (0.0%) | 4/147 (2.7%) | 0/67 (0.0%) | 0/31 (0.0%) | 0/10 (0.0%) | 9/331 (2.7%) |
| MYCN amplification | 0/47 (0.0%) | 1/11 (9.1%) | 7/118 (5.9%) | 13/55 (23.6%) | 0/27 (0.0%) | 1/5 (20.0%) | 22/263 (8.4%) |
| MYC amplification | 10/47 (21.3%) | 1/11 (9.1%) | 2/118 (1.7%) | 0/54 (0.0%) | 0/27 (0.0%) | 0/5 (0.0%) | 13/262 (5.0%) |
| TERT | 0/42 (0.0%) | 0/10 (0.0%) | 1/108 (0.9%) | 16/51 (31.4%) | 0/25 (0.0%) | 1/6 (16.7%) | 18/242 (7.4%) |
| TP53 | 1/63 (1.6%) | 0/12 (0.0%) | 1/143 (0.7%) | 12/67 (17.9%) | 1/31 (3.2%) | 0/8 (0.0%) | 15/324 (4.6%) |
| CTNNB1 | 0/50 (0.0%) | 0/11 (0.0%) | 1/109 (0.9%) | 0/53 (0.0%) | 25/28 (89.3%) | 0/6 (0.0%) | 26/257 (10.1%) |
| **MB(Grp3/Grp4) methylation subtype** | | | | | | | |
| I | 2/63 (3.2%) | 1/13 (7.7%) | 2/147 (1.4%) | 0/67 (0.0%) | 0/31 (0.0%) | 0/10 (0.0%) | 5/331 (1.5%) |
| II | 22/63 (34.9%) | 0/13 (0.0%) | 0/147 (0.0%) | 0/67 (0.0%) | 0/31 (0.0%) | 0/10 (0.0%) | 22/331 (6.6%) |
| III | 10/63 (15.9%) | 4/13 (30.8%) | 0/147 (0.0%) | 0/67 (0.0%) | 0/31 (0.0%) | 1/10 (10.0%) | 15/331 (4.5%) |
| IV | 16/63 (25.4%) | 1/13 (7.7%) | 0/147 (0.0%) | 0/67 (0.0%) | 0/31 (0.0%) | 0/10 (0.0%) | 17/331 (5.1%) |
| V | 3/63 (4.8%) | 2/13 (15.4%) | 11/147 (7.5%) | 0/67 (0.0%) | 0/31 (0.0%) | 1/10 (10.0%) | 17/331 (5.1%) |
| VI | 0/63 (0.0%) | 1/13 (7.7%) | 18/147 (12.2%) | 0/67 (0.0%) | 0/31 (0.0%) | 0/10 (0.0%) | 19/331 (5.7%) |
| VII | 1/63 (1.6%) | 1/13 (7.7%) | 35/147 (23.8%) | 0/67 (0.0%) | 0/31 (0.0%) | 0/10 (0.0%) | 37/331 (11.2%) |
| VIII | 0/63 (0.0%) | 0/13 (0.0%) | 48/147 (32.7%) | 0/67 (0.0%) | 0/31 (0.0%) | 0/10 (0.0%) | 48/331 (14.5%) |
| Unknown/NA | 9/63 (14.3%) | 3/13 (23.1%) | 33/147 (22.4%) | 67/67 (100.0%) | 31/31 (100.0%) | 8/10 (80.0%) | 151/331 (45.6%) |
| **Methylation subgroup** | | | | | | | |
| Group3 | 52/63 (82.5%) | 6/13 (46.2%) | 3/147 (2.0%) | 0/67 (0.0%) | 0/31 (0.0%) | 1/10 (10.0%) | 62/331 (18.7%) |
| Group4 | 6/63 (9.5%) | 5/13 (38.5%) | 125/147 (85.0%) | 0/67 (0.0%) | 0/31 (0.0%) | 1/10 (10.0%) | 137/331 (41.4%) |
| SHH | 0/63 (0.0%) | 0/13 (0.0%) | 0/147 (0.0%) | 0/67 (0.0%) | 29/31 (93.5%) | 0/10 (0.0%) | 29/331 (8.8%) |
| WNT | 0/63 (0.0%) | 0/13 (0.0%) | 0/147 (0.0%) | 61/67 (91.0%) | 0/31 (0.0%) | 6/10 (60.0%) | 67/331 (20.2%) |
| Unknown/NA | 5/63 (7.9%) | 2/13 (15.4%) | 19/147 (12.9%) | 6/67 (9.0%) | 2/31 (6.5%) | 2/10 (20.0%) | 36/331 (10.9%) |
